# Post-COVID impairment of memory T cell responses to community-acquired pathogens can be rectified by activating cellular metabolism

**DOI:** 10.64898/2025.12.31.697156

**Authors:** Daniel D. Carroll, Kamacay Cira, Jack Archer, Jason Shapiro, Ue-Yu Pen, David Tieri, Lucia Leonor, Neda D. Roofchayee, Samantha S. Yee, Marc Wahab, Igor J. Koralnik, John C. Alverdy, Arjun S. Raman, Lavanya Visvabharathy

## Abstract

Infection rates involving bacterial and viral pathogens have increased precipitously after the COVID-19 pandemic. While it has been speculated that higher infection rates resulted from increased hospitalizations throughout the pandemic or greater use of antibiotics, precisely why rates remain high today has remained unexplained. Mitochondrial dysfunction is known to occur post-COVID and may disrupt immune responses. Within T cells, SARS-CoV-2 infection is linked to low mitochondrial membrane potential, increased mitochondrial apoptosis, and decreased mitochondrial respiration, which together impact cellular activation and function beyond the acute phase of illness. Here, we demonstrate that decreased mitochondrial function in antigen-specific T cells post-COVID may contribute to higher infection susceptibility by metabolically immobilizing T cell memory responses. Using donor-matched peripheral blood samples from 31 COVID-naïve individuals who subsequently contracted COVID-19, we tracked how influenza A (IAV), *Staphylococcus aureus* (SA), and Varicella-zoster virus (VZV) T cell responses were impacted by COVID-19 infection. We found that gene expression linked to T cell activation decreased but mitochondrial redox pathways increased in CD4 memory T cells post-COVID. However, mitochondrial flux and reactive oxygen species production were limited in a plurality of post-COVID memory T cells after stimulation with IAV, SA, and VZV. Furthermore, we found a disordered relationship between memory T cell mobilization of glycolysis, fatty acid metabolism, and oxidative phosphorylation pathways post-COVID which resulted in diminished use of catabolic pathways including glycolysis and fatty acid oxidation in antigen-specific T cells. Modulation of mitochondrial function with metformin and ubiquinol partially rescued the post-COVID decline in T cell catabolism. Collectively, these findings indicate that COVID-19 infection may have lasting effects on inhibiting T cell memory responses to commonly encountered community-acquired pathogens which can be corrected with commonly available medications. This has significant implications for the clinical care of immunologically vulnerable populations in the post-pandemic era.

## Introduction

Infections of all types have increased worldwide after the COVID-19 pandemic. Over the past 3 years (2022-24), there have been outbreaks of viral Mpox^1^, co-occurring epidemics of respiratory syncytial virus (RSV), influenza, and COVID-19^2^, as well as increased rates of bacterial infections in hospitalized patients above pre-pandemic levels^3^. Co-occurring epidemics, particularly with pathogens that occupy the same host niche, were previously considered rare. Epidemiological studies on respiratory viral infection cases from 2005-13 found negative temporal correlations between influenza virus, RSV, and other common cold virus rates, demonstrating an ecological “competitive inhibition” when multiple pathogens target the same host niche^4^. However, this pattern has been upended by COVID-19. Frequent observations of co-occurring respiratory virus outbreaks^5^ as well as elevated herpes virus reactivation^6^ and *Staphylococcus aureus* infections^7^ occur during and proximal to a COVID-19 infection. Explanations put forth have ranged from increased hospital usage during the COVID-19 pandemic^8^ to greater use of antibiotics^9^, but none of these explain why infection rates continue to increase in the present day. We propose an alternate hypothesis: that COVID-19 may affect T cell memory responses to other pathogens by reprogramming mitochondrial metabolism.

Mitochondrial dysfunction is a hallmark of acute and post-acute COVID-19. Patients hospitalized for COVID-19 had unique T cell populations characterized by downregulated glycolytic proteins and linked to lymphocyte apoptosis and increased disease severity^10^. Importantly, this phenotype was unique to COVID-19 patients and not observed in those with hepatitis C or those hospitalized for influenza. Similarly, studies have observed increased T cell mitochondrial mass, altered mitochondrial reactive oxygen species (ROS) levels, metabolic quiescence, and disrupted mitochondrial architecture in acute COVID-19 patients^11^. Importantly, mitochondrial regulation of T cell metabolism is critical to maintain a balance between effector and memory functions through preferential utilization of glycolysis or OXPHOS as a means to generate energy^12^. At baseline, resting memory T cells have a metabolically quiescent phenotype characterized by lower mitochondrial membrane potential and increased mitochondrial biogenesis, which are desirable features of long-lived cells that protect against reinfection. Upon antigen re-encounter, conversely, memory T cells exploit their higher spare respiratory capacity (SRC) relative to effector cells to provide the ATP necessary through rapid mobilization of glycolysis to differentiate into secondary T effector cells^13^. This process is crucial to mediate the rapid recall ability of memory T cells. However, whether and how SARS-CoV-2 infection impacts the critical switch between glycolysis and OXPHOS in memory T cells to induce long-term immunosuppression remains an open question.

Here, we demonstrate that antigen-specific memory T cells from post-COVID individuals show lower glycolysis usage and disordered mitochondrial metabolism in response to commonly encountered childhood and environmental pathogens. Critically, we studied samples from individuals who were initially COVID-naïve, i.e. they had never been exposed to SARS-CoV-2 at the time of blood collection. Using matched samples from the same individuals, we compared T cell memory responses in pre-COVID and post-COVID peripheral blood mononuclear cells (PBMCs) to Influenza A virus (IAV), *Staphylococcus aureus* (SA), and Varicella-zoster virus (VZV). Despite unchanged antigen-specific memory T cell percentages, post-COVID memory T cell responses exhibited reduced coordination across glycolysis, fatty-acid synthesis/oxidation, and OXPHOS pathways relative to pre-COVID responses, a pattern associated with higher PD1 expression.. This deficit was partially rescued by exposure to pharmacological agents that activate mitochondrial complex I and III, demonstrating potential novel avenues for treatment of post-COVID immunosuppression. Overall, we uncover a COVID-related deficit in T cell memory responses to community-acquired pathogens which may explain the durability of elevated infection rates post-pandemic.

## 3 Results

### Increase in infections and associated mortality rates post-pandemic

Several studies have reported increased rates of both bacterial and viral infections post-pandemic which prompted us to investigate whether similar trends were evident in publicly available nationwide infection data from the Centers for Disease Control and Prevention (CDC). The CDC WONDER (Wide-ranging Online Data for Epidemiologic Research) database is an online tool to output publicly available data from the CDC pertaining to morbidity, mortality, and other public health metrics from across the US from 1999-2023^14^. Data can be parsed by disease type and locale, among other metrics. We limited our search to the 3 years immediately preceding the pandemic (2017-2019) compared to 3 years after the pandemic began (2021-2023), excluding 2020 due to disruptions in data collection which arose with COVID-19. Nationwide, death rates associated with bacterial infections significantly increased post-2020, with particular increases in *Staphylococcus* and Group A Streptococcus (GAS) deaths per 100,000 compared to pre-pandemic years (Fig. 1A). Ventilator associated infections which frequently consist of respiratory viral infections were similarly increased (Fig. 1A). Illinois state level data, where our cohort is based, had similar trends showing a significant upsurge in hospital-acquired bloodstream infections (CLABSI) and *Staphylococcal*-associated deaths (Fig. 1B). This led us to investigate the associated between SARS-CoV-2 infection and immune memory to other common pathogens.

**Figure 1:**
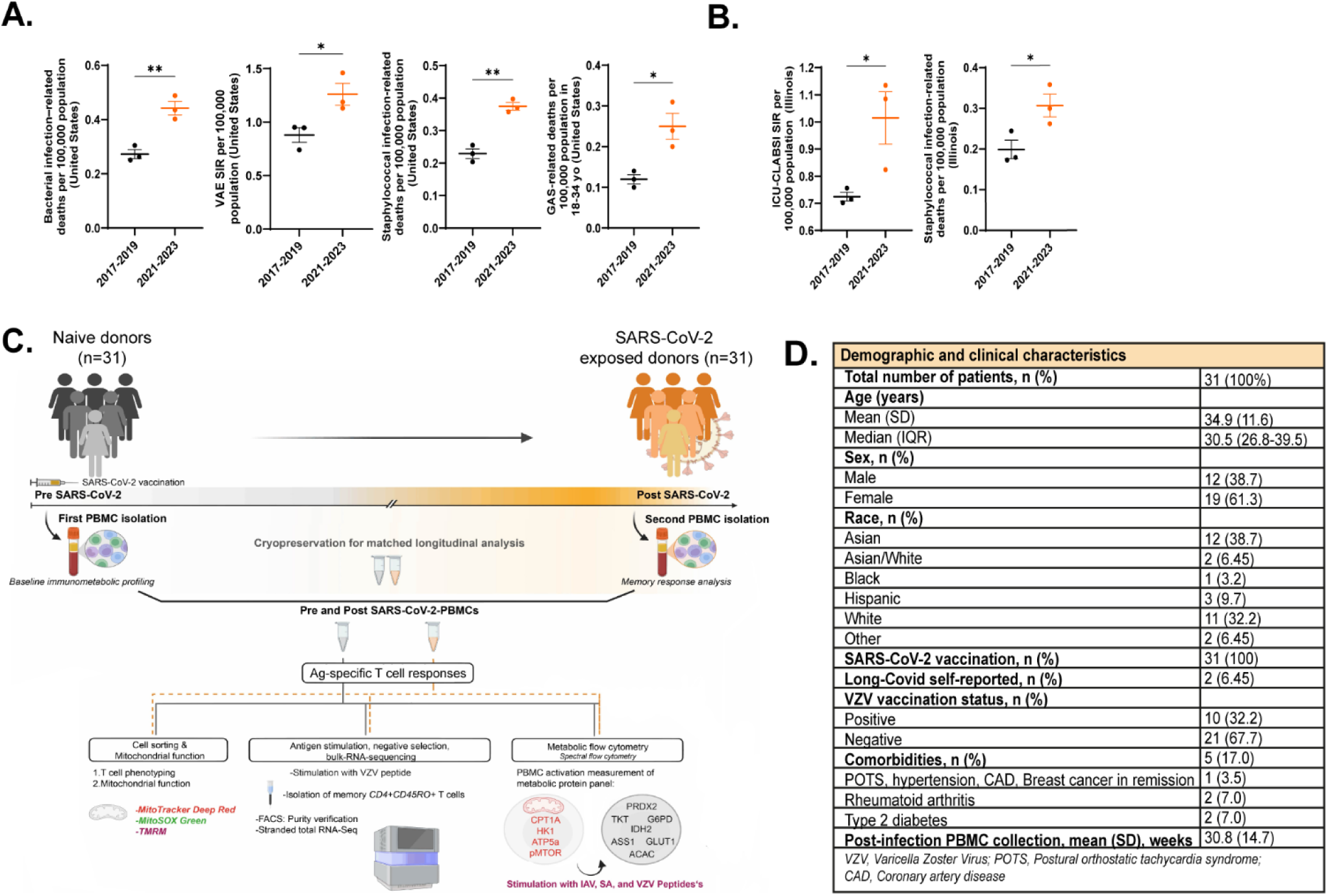
Infection rates post-COVID pandemic, study design, demographic information. A.) Bacterial infection-related deaths, ventilator-associated respiratory infections, *Staphylococcal* infection-related deaths, and Group A Streptococcus (GAS)-related deaths per 100,000 significantly increased in the US during the years after the COVID-19 pandemic. B.) Total infection-related deaths, bloodstream infection rates, and *Staphylococcal* infection-related deaths similarly increased in the state of Illinois after the COVID-19 pandemic. C.) Study design schematic. D.) Study participant demographics. VAE SIR: ventilator-associated events, standardized infection rate. ICU-CLABSI: intensive care unit-central line-associated bloodstream infections. *p<0.05, **p<0.01, ***p<0.005 by unpaired two-tailed t-test. Error bars show mean ± SEM. Data for A-B gathered from the CDC Wonder database in September 2024.

### Mitochondrial oxidative phosphorylation and antigen presentation pathway gene expression in memory T cells is significantly impacted by COVID status

Our group had collected peripheral blood mononuclear cell (PBMC) samples from 31 healthy, COVID-naive donors from 2021-2023 as part of the IRB-STU00212583 study at Northwestern University investigating T cell responses to COVID-19 vaccination. COVID-naïve status was confirmed by lack of antibody responses to SARS-CoV-2 nucleocapsid protein prior to enrollment (Fig. S1A). During the course of the study (2021-24), each participant’s COVID status was confirmed by either PCR– or antigen test for SARS-CoV-2. As we collected samples longitudinally to track vaccine-elicited immunity, we were able to cryobank matched samples from the same individuals pre– and approximately 30 weeks post-COVID infection. One patient contracted COVID twice during the course of the study, and we used samples from all 3 timepoints in the analysis. Given the increased infection rates and sustained cell and mitochondrial dysfunction post-COVID^15,16^, we used these matched samples to formally test the hypothesis that SARS-CoV-2 increases susceptibility to other infections by altering memory T cell metabolic programs (study design in Fig. 1C, demographics in Fig. 1D).

To determine the impact of COVID-19 on T cell memory, we initially chose to study Varicella-zoster virus (VZV) responses. We chose VZV for the following reasons. First, all individuals would have exposure from either natural infection or vaccination in childhood. In addition, VZV-specific recall responses remain stable over periods of 24 months or more and persist at high levels into adulthood, making it unlikely for VZV responses to change across the sampling interval^17^. Reports have also credibly shown that increased VZV reactivation is found in COVID-19 survivors^18^. Pre-COVID samples had a 93% VZV antibody positivity rate (Fig. S1B), indicating that the majority of study participants had humoral immune memory to VZV. This established a uniform baseline of exposure against which to interrogate post-COVID changes in recall quality. Therefore, sixteen subjects were randomly selected from the original cohort of thirty, and their PBMCs were stimulated with a VZV Orf4 peptide array spanning the entire protein (antigen descriptions in Fig. S1C). We then purified CD4+CD45RO+ memory T cells by negative selection (purity in Fig. S2) before isolating RNA. Further analysis was conducted with bulk RNA sequencing, which provided us with a transcriptional signature of pre– and post-COVID T-cell modifications (Fig. 2A).

**Figure 2:**
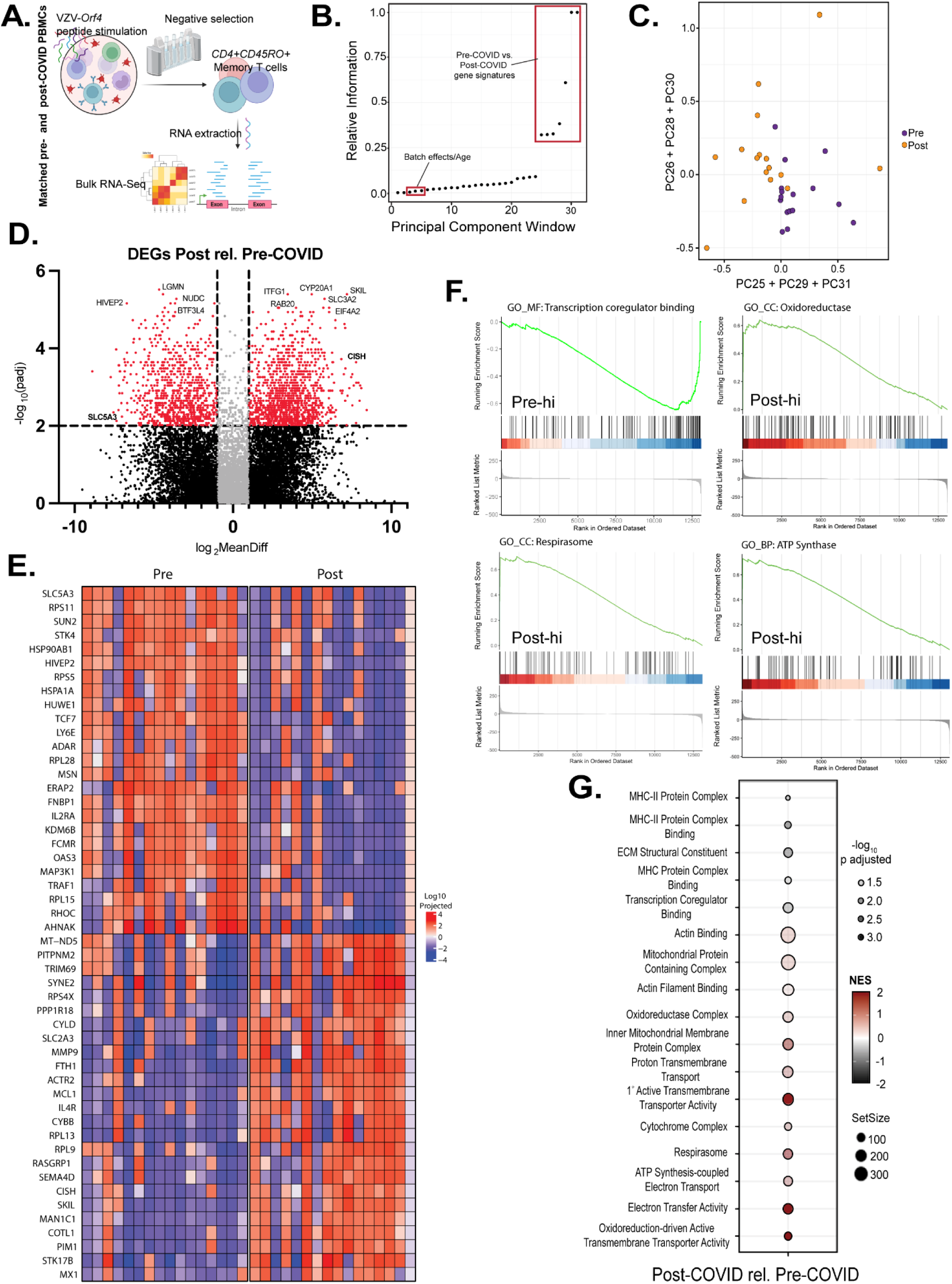
Enhanced mitochondrial OXPHOS and diminished MHC-II antigen presentation gene expression in post-COVID CD4 memory T cells after VZV antigen stimulation. A.) Experimental design. B.) Statistical information about variance in pre– vs. post-COVID gene signatures are contained in principal components 25-31. C.) Isolating principal components 25-31 effectively separates samples based on pre– vs. post-COVID gene signatures. D.) Differential gene expression hits in pre– vs. post-COVID CD4 memory T cells. E.) Top 25 differentially expressed genes in pre– vs. post-COVID samples. F.) Gene set enrichment analysis (GSEA) shows significantly decreased transcription factor binding genes and elevated ATP synthase, oxidoreductase, and mitochondrial respirasome pathway genes in post-COVID CD4 memory T cells. Pre-hi: pathway expression higher in pre-COVID samples. Post-hi: pathway expression higher in post-COVID samples. G.) Significantly altered gene expression pathways enriched in mitochondrial function-related cellular processes in post-COVID relative to pre-COVID CD4 memory T cells. DEGs were calculated based on p_adj_=0.05 using a Bonferroni correction for multiple comparisons. Log_2_MeanDiff= mean of gene expression difference, log 2. N=16 matched pre– and post-COVID samples.

To reveal differences between these two cell populations, we turned to dimension-reduction approaches typically used in analyzing high-content biological data^19^. We first performed Principal Components Analysis (PCA), defining a set of 31 principal components (PCs). In general, PCA focuses signals of statistical variation amongst the so-called ‘top-modes’ of variance—modes that harbor the most amount of variation within the data. Analysis of the top PCs in our data illustrated a statistically significant, non-random relationship with batch effect and age of the patient from which the sample was collected (Fig. S3A). Further analysis using non-linear dimension-reduction methods—UMAP and t-SNE—recapitulated the same finding (Fig. S3B). These results suggested that the most global signals of variation in our data were unrelated to pre-versus post-COVID signatures of T cell modification.

A parsimonious explanation of our findings would suggest that there may not exist a transcriptional signature separating pre-from post-COVID T cells. However, recently created statistical approaches, developed within the field of bacterial genomics and phylogenetic analysis, have suggested that relevant biological signal may actually reside in PCs that harbor substantially less variation than the top PCs. Fundamentally, this is because the spectrum of principal components encodes a tree of relatedness amongst systems, where biological information captured within the lowest PCs is statistically nested within the information captured by the highest, most dominant PCs.

With this nested data structure in mind, we next analyzed our data using a framework called SCALES (Spectral Correlation Analysis of Layered Evolutionary Signals)—a statistical approach that performs a data-driven search across all PCs to identify regions of the PC spectrum most associated with the desired phenotype^19^. (Fig. 2B). Application of SCALES identified PCs encoding information regarding a “COVID effect” and revealed distinct transcriptional signatures separating pre– and post-COVID states (Fig. 2B,C). Differential expression analysis identified extensive transcriptional remodeling, with 2048 differentially expressed genes (DEGs) between pre– and post-COVID samples (p_adj_ < 0.05, log_2_MeanDiff >1; Fig. 2D), where log₂MeanDiff denotes the log₂ difference between the mean projected expression in post-versus pre-COVID samples along the selected principal components. Transcripts most strongly downregulated post-COVID included solute carrier family 5 member 3 (*SLC5A3*), transcription factor 7 (*TCF7*), lymphocyte antigen 6 family member E (*LY6E*), endoplasmic reticulum aminopeptidase 2 (*ERAP2*), and interleukin-2 receptor subunit alpha (*IL2RA*), among others. These genes are involved in inositol import and calcium mobilization for T cell activation (*SLC5A3*), memory T-cell differentiation^20^ (*TCF7*), coronavirus entry restriction factor^21^ (*LY6E*), and cytokine-driven proliferation (*IL2RA*). In contrast, post-COVID samples were enriched for MX dynamin-like GTPase 1 which may promote viral persistence^22^ (*MX1*), cytokine-inducible SH2-containing protein which suppresses T cell activation^23^ (*CISH*), and mitochondrially encoded NADH dehydrogenase 5 (*MT-ND5*), among others (Fig. 2E). Importantly, pathway analysis found significant downregulations in MHC-II complex binding and transcription factor co-regulation, but upregulated mitochondrial OXPHOS and ATP synthesis pathways in post-COVID T cells (Fig. 2F, G). Together, these results demonstrate that SARS-CoV-2 infection is accompanied by durable transcriptional reprogramming of VZV antigen-stimulated CD4⁺ memory T cells, defined by enrichment of genes pertinent to mitochondrial metabolism. This is consistent with a metabolic shift that may contribute to the functional impairments of CD4⁺ memory T cells within this cohort.

### Bimodal phenotype of mitochondrial function in antigen-specific memory T cells post-COVID

Because post-COVID memory T cells show enrichment of mitochondrial-metabolic pathways, we inferred that these transcriptional shifts indicate functional metabolic changes that may impair antigen-specific memory responses. To test this, we expanded our analyses beyond VZV to include Influenza A virus (IAV) and *Staphylococcus aureus* (SA) antigens representing exposure of broader clinical relevance. All participants showed prior exposure to these pathogens, confirmed serologically (Fig. S1D).

We first assessed mitochondrial activity at baseline using functional dyes: Mitotracker (MTT) to detect active mitochondrial mass, tetramethylrhodamine methyl ester (TMRM) to assess mitochondrial membrane potential and electron flux, and MitoSOX (SOX) to detect mitochondrial reactive oxygen species (ROS) production. Baseline values were comparable across unstimulated CD3, CD4, CD8 T cells and CD14 monocytes pre– and post-COVID (Fig. 3A; gating in Fig. S4). Similarly, the overall frequencies of antigen-specific activation-induced marker (AIM+; gating in Fig. S5) CD4⁺ and CD8⁺ memory T cells recognizing VZV, SA, or IAV antigens (described in Fig. S1C) were preserved, apart from a modest increase in VZV-specific AIM+ CD8⁺ memory T cells post-COVID (Fig. 3B, E, H). These results indicate that SARS-CoV-2 infection does not substantially alter the pool size of antigen-specific memory T cells.

**Figure 3:**
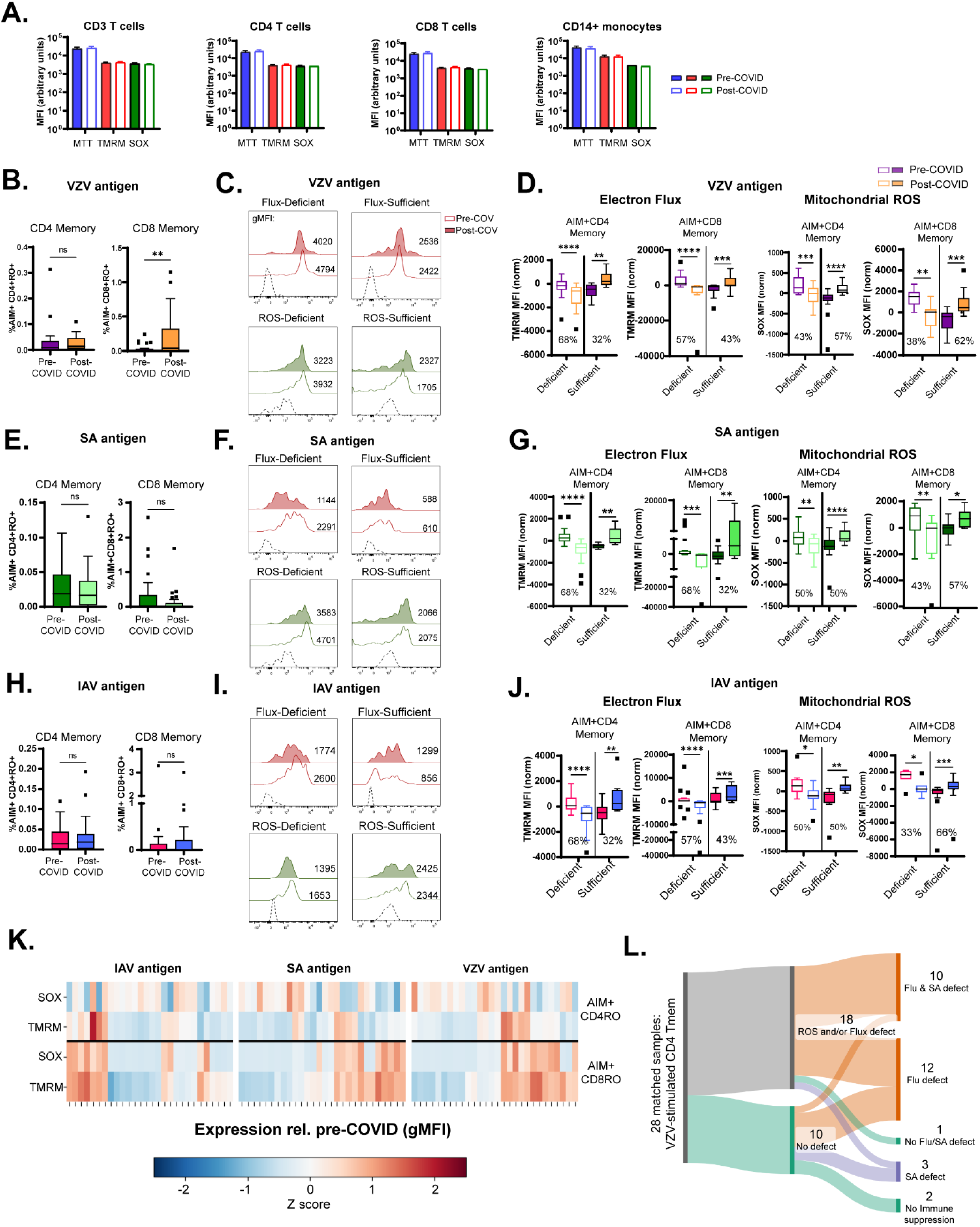
Bimodal phenotype of mitochondrial dysfunction in antigen-stimulated memory T cells post-COVID. A.) Baseline levels of active mitochondrial mass (MTT), mitochondrial electron flux (TMRM), and mitochondrial reactive oxygen species production (SOX) in immune cell subsets are unchanged pre– and post-COVID. B.) VZV antigen-specific (AIM+) CD4 memory T cell numbers are unchanged after COVID, while AIM+ CD8 memory T cell numbers are elevated. C-D.) Bimodal phenotype of mitochondrial ROS and mitochondrial flux post-COVID in VZV antigen-specific memory T cells. A majority of study subjects produced less mitochondrial ROS and displayed lower electron flux post-COVID. E.) Levels of SA antigen-specific memory T cells are unchanged post-COVID. F-G.) Deficiency in electron flux and mitochondrial ROS production in SA antigen-specific memory T cells in a majority of subjects. H.) Unchanged percentages of IAV antigen-specific memory T cells post-COVID. I-J.) Similar to VZV and SA, bimodal phenotype of mitochondrial ROS and mitochondrial flux is evident post-COVID in IAV antigen-specific memory T cells. K.) Heatmap of mitochondrial ROS and mitochondrial flux expression across all samples. L.) Flow diagram showing that 92.8% of subjects had ROS and/or flux deficiency to at least one set of antigens. Graphs show mean ±SEM. *p<0.05, **p<0.01, ***p<0.005, ****p<0.0001 by paired Student’s t test or Wilcoxon test from n=28 individuals, matched samples. C, F, I: representative data from 1/28 individuals; dashed histogram represents FMO.

Functional analyses, however, revealed post-COVID impairments in mitochondrial responses upon antigen re-exposure in a bimodal fashion. VZV-specific CD4⁺ and CD8⁺ memory T cells from more than half of the study participants had significantly lower mitochondrial flux and ROS production post-COVID compared to paired pre-COVID samples (Fig. 3C, D). Comparable patterns were observed following stimulation with SA (Fig. 3F, G) and IAV (Fig. 3I, J). No significant differences were found in mitochondrial mass when parsed by pre– vs. post-COVID status, with the exception of SA-stimulated CD4 memory T cells (Fig. S6). The population distribution of TMRM and SOX expression in memory T cells is summarized as a heatmap in Figure 3K. Across all antigen stimulations, 33-79% of individuals displayed post-COVID deficits in at least one mitochondrial function parameter. A participant-level flow diagram confirmed the breadth of these defects: 93% of subjects exhibited impaired mitochondrial function in antigen-specific CD4⁺ memory T cells for at least one pathogen, and most showed deficits across multiple pathogens (Fig. 3L). Together, these findings demonstrate that although antigen-specific memory frequencies remain stable, SARS-CoV-2 infection is associated with broad, pathogen-independent impairments in mitochondrial bioenergetics underpinning T cell recall responses.

### Limited connectivity of metabolic pathways in post-COVID memory T cells

Given the disruption of mitochondrial function in antigen-specific memory T cells following COVID-19 infection, we further investigated how specific cellular metabolic pathways were affected. We employed the Met-Flow technique^24^ to define metabolic state changes in immune cells after activation. Met-Flow uses high-dimensional spectral flow cytometry to detect rate-limiting enzymes involved in glycolysis, oxidative phosphorylation, and fatty acid oxidation/ synthesis. This technique can characterize and quantify metabolic alterations associated with T cell survival, differentiation, activation, and function at the single cell level, allowing for subset-specific data collection. We focused on the following enzyme targets: GLUT1, HK1, G6PD, TKT, PRDX2, ASS1, ACAC, IDH2, CPT1A, and ATP5a (diagram and enzyme functions: Fig. 4A) involved in glycolysis, fatty acid metabolism, arginine metabolism, and OXPHOS.

**Figure 4:**
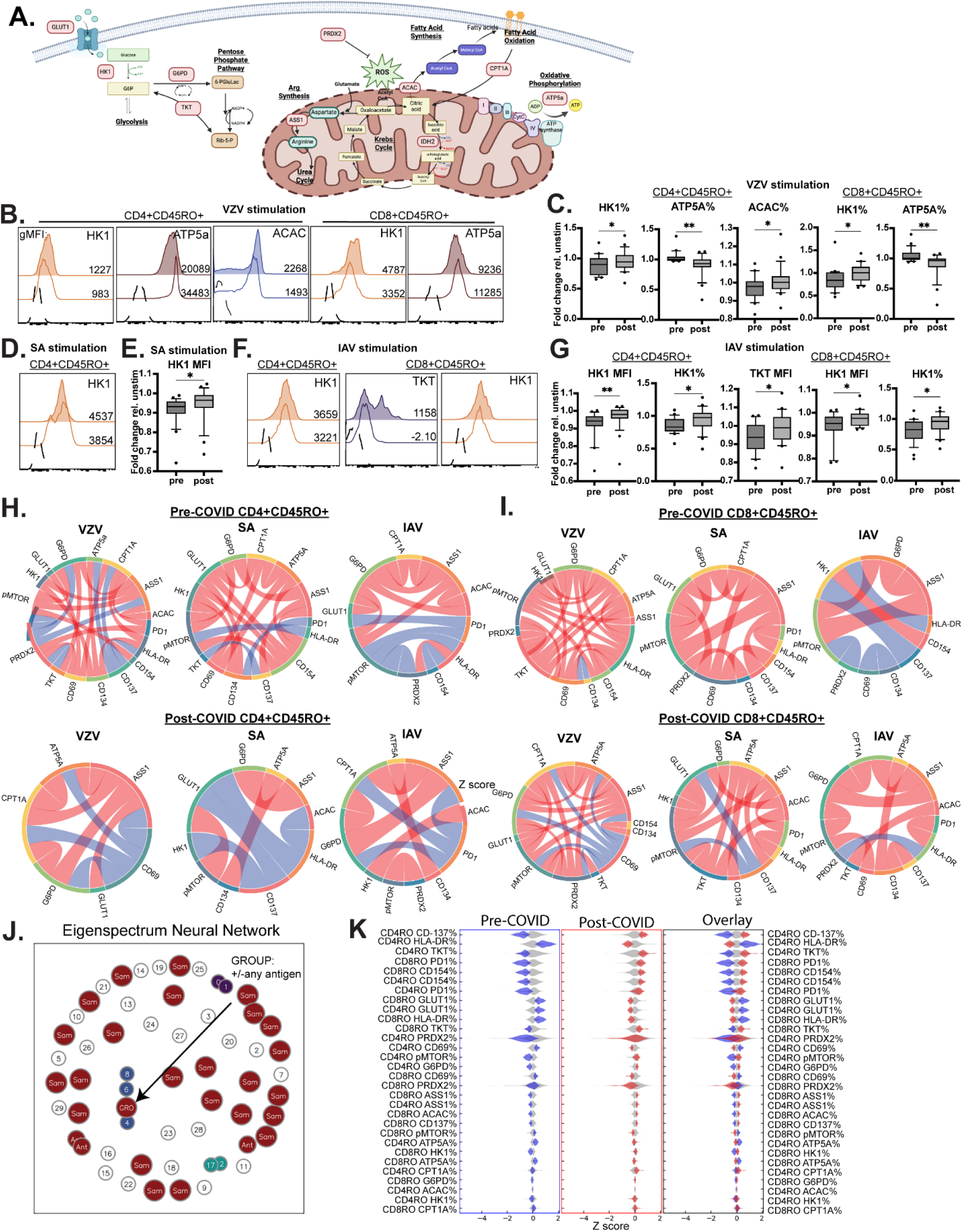
Metabolic analysis at the cell population level reveals post-COVID elevations in glycolytic enzymes but diminished OXPHOS enzyme levels in memory T cells. A.) Metabolic enzyme targets quantified in flow cytometry (MetFlow). Specific targets are in red boxes. B-C.) Post-COVID elevations in HK1 (glycolysis) and ACAC (fatty acid synthesis) and decreased expression of ATP5a (OXPHOS) in memory T cells after VZV antigen stimulation. D-E.) Higher HK1 expression in SA antigen-stimulated CD4 memory T cells post-COVID. F-G). Elevated HK1 expression and TKT expression (both glycolysis enzymes) in IAV-stimulated memory T cells after COVID. H-I.) Connectivity between metabolic pathways (% enzyme or activation marker positive memory T cells) is lost post-COVID in both CD4 (H) and CD8 (I) memory T cells. The loss of connectivity is more pronounced in CD4 memory T cells. Blue arcs: positive correlation; red arcs: negative correlation. J.) Eigenspectrum neural network analysis of statistical connections within CD4 and CD8 memory T cells finds a group-specific pre– vs. post-COVID signature. K.) Feature distribution map differentiates pre– vs. post-COVID memory T cell phenotypes. Blue: post-COVID; red: pre-COVID. The top 13 features explain the majority of statistical variation between pre– and post-COVID groups (overlay). Filled histograms: post-COVID; open histograms: pre-COVID. Graphs show mean ±SEM. *p<0.05, **p<0.01 by paired Student’s t test in n=24 samples. H, I: chord diagrams of Spearman correlations in % of memory T cells positive for the specified marker. J, K: data from B-I used to generate feature maps using Boltzmann Brain analysis. Feature maps show clear segregation of biomarkers differentiating pre-COVID (CD137^lo^, HLA-DR^hi^, PD1^lo^, GLUT1^hi^) from post-COVID (CD137^hi^, HLA-DR^lo^, PD1^hi^, GLUT1^lo^) T cell states.

At baseline, there were few differences in CD4 and CD8 memory T cell expression of key metabolic enzymes post-COVID, apart from elevated HK1 levels in CD8 memory cells (Fig. S7). However, VZV antigen stimulation led to higher HK1 and ACAC expression but lower ATP5a expression in post-COVID memory T cells compared to matched pre-COVID controls (Fig. 4B-C, all data show fold change relative to unstimulated samples). Similarly elevated HK1 expression as well as TKT expression were found after SA (Fig. 4D-E) and IAV antigen stimulation (Fig. 4F-G), demonstrating that usage of glycolysis pathways in memory T cells was elevated but OXPHOS was lower at the CD4 and CD8 memory T cell population level after COVID. However, a different picture emerged when examining how metabolic pathways were connected within cells. Post-COVID memory T cells displayed markedly diminished connectivity between metabolic enzyme expression and key activation markers such as CD69, HLA-DR, and CD137 compared to matched pre-COVID controls (Fig. 4H, I).

To better define data features underlying the lack of metabolic connectivity in memory T cells, we sought to create a statistical embedding that differentiated pre-versus post-COVID pathways. To do this, we employed an analysis program called the Boltzmann Brain—a program that constructs embeddings from the eigenspectrum of data. While detailed in Methods, briefly this program conducts a data-driven search for embeddings that are maximally information-rich with respect to a user-defined phenotype of interest. The embeddings can then be queried to discover features that are a statistically inferred ‘biomarker’ of phenotype. In our case, Boltzmann analysis revealed a unique relationship between “Group” (pre– or post-COVID) and “Antigen” (VZV/SA/IAV stimulation) when evaluating memory T cell expression of metabolic and activation markers (Fig. 4J), the same underlying data that we used to generate Figures 4B-I. We analyzed the feature distribution characterizing antigen stimulation of memory T cells and found distinct biomarker signatures differentiating samples by pre– vs. post-COVID status (Fig. 4K). The top features responsible for the largest differences showed elevated HLA-DR and GLUT1 in pre-COVID samples but enriched CD137, TKT, and PD1 expression in post-COVID samples. These data demonstrate that post-COVID memory responses are characterized by diminished connectivity between metabolic pathways but higher glycolysis usage and expression of the inhibitory marker PD1 after antigen stimulation at the cell population level.

### Antigen-specific memory T cells show deficiencies in glycolysis/fatty acid oxidation usage post-COVID, which is partially corrected with mitochondrial activator drugs

Though memory T cell responses at the population level showed increased glycolysis usage as shown by elevated HK1 expression (Fig. 4B-C), this did not necessarily reflect the behavior of antigen-specific memory T cells. To address this, we assessed the response of AIM+ CD4 and CD8 memory T cells to IAV, SA, and VZV stimulation. In contrast to phenotypes at the cell population level, AIM+ memory T cells displayed significantly *lower* HK1 and CPT1A expression (representing glycolysis and FAO metabolic pathways) but higher ACAC and pMTOR expression (representing FAS and cell proliferation pathways) indicating mobilization of anabolic pathways, but diminished use of catabolic glycolysis and fatty acid oxidation pathways after antigen stimulation (Fig. 5A-B).

**Figure 5:**
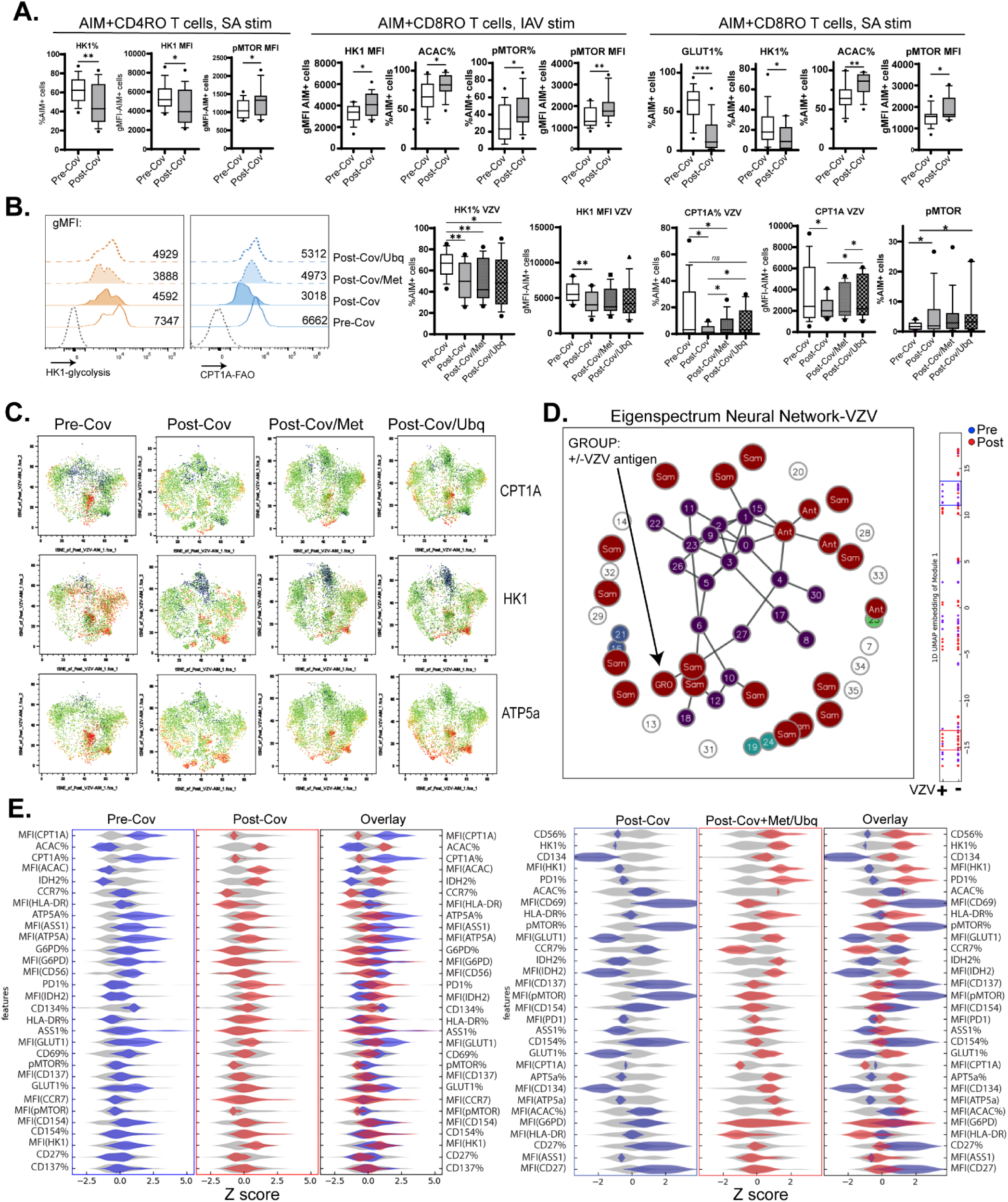
Metabolic analysis of antigen-specific memory T cells shows lower glycolysis usage and alterations in fatty acid metabolism post-COVID, which is partly reversed by pharmacological activation of mitochondrial function. A.) Decreased glycolysis (GLUT1, HK1) but elevated fatty acid synthesis (ACAC) and phosphor-mTOR in SA and IAV antigen-specific memory T cells post-COVID. B.) Deficits in glycolysis and fatty acid oxidation in VZV antigen-specific CD4 memory T cells can be partly rescued by modulation of mitochondrial complex I (Met, dotted filled histogram) or complex III (Ubq, dotted open histogram). C.) Expression of CPT1A, HK1, and ATP5a in VZV-specific CD4 memory T cells is partly rescued by exposure to Met or Ubq. Data are representative of n=16 samples. D.) Eigenspectrum neural network of VZV antigen-specific CD4 T cells after discovery of a pre– vs. post-COVID (“Group”) MetFlow signature. Bar to the right represents 1D UMAP representation of data where blue dots are pre-COVID samples and red dots are post-COVID samples. Red and blue windows represent areas with the greatest feature divergence between pre– and post-COVID VZV-specific CD4 memory T cells. Feature maps in 5E were derived from information contained within these windows. E.) *Left:* Feature distribution map of pre– vs. post-COVID VZV-specific CD4 memory T cells showing higher glycolysis (GLUT1, HK1), lower immunosuppression (pMTOR), and higher fatty acid oxidation (CPT1A) in pre-COVID samples relative to post-COVID samples. *Right:* Feature distribution map of post-COVID VZV-specific T cells treated with Met or Ubq shows higher expression of glycolytic enzymes (HK1), activation markers (CD134, HLA-DR), and lower pMTOR expression compared to untreated T cells. Graphs show mean ±SEM. *p<0.05, **p<0.01 by paired Student’s t test in n=16 samples.

The switch from OXPHOS to glycolysis is essential for memory T cells to quickly respond to pathogen re-encounter. Similarly, memory T cells rely on fatty acid oxidation to activate effector programs^25^. Given that post-COVID antigen-specific memory T cells were deficient in expression of the glycolytic enzyme HK1 and FAO enzyme CPT1A, we determined whether pharmacological modulation of mitochondrial function could rescue post-COVID deficits in cellular metabolism. We focused on CD4 memory responses to VZV because of transcriptional (Fig. 2) and mitochondrial function (Fig. 3) signatures were able to discriminate between pre– and post-COVID groups. Metformin and ubiquinol are two pharmacological modifiers of mitochondrial metabolism with distinct physiological activities. Metformin is a commonly prescribed anti-diabetic drug that can lower blood glucose levels by promoting glycolysis/FAO and decreasing mitochondrial complex I activation^26^. It has also been shown to promote memory T cell responses against infections and tumors ^27,28^. Ubiquinol (also known as coenzyme Q10) is a potent anti-oxidant that limits inflammation and activates mitochondrial complex III which can promote T cell memory generation to infections and cancer^29^.

Incubation of PBMCs with 10µM metformin or 5µM ubiquinol had a demonstrable effect in enhancing HK1 and CPT1A expression in VZV-specific CD4 memory T cells (Fig. 5B), though not to pre-COVID levels. VZV-specific T cells showed a marked enhancement in CPT1A, HK1, and ATP5a expression after drug treatment, with ubiquinol being more potent than metformin in promoting expression (Fig. 5C). After analysis with the Boltzmann Brain program (Fig. 5D), we identified several data features that distinguished pre– and post-COVID VZV-specific memory T cells and unique signatures of post-COVID T cells after drug treatment. Pre-COVID VZV-specific T cells displayed elevated glucose import (higher GLUT1 expression), costimulation/activation (CD134 expression), and lower fatty acid synthesis (ACAC expression) in addition to elevated glycolysis/FAO (HK1/CPT1A expression; Fig. 5E, left panel). Metformin or ubiquinol treatment was able to partially restore the post-COVID deficits in HK1 expression, CD134 expression, and GLUT1 expression (Fig. 5E, right panel), demonstrating that commonly available pharmacological modulators of mitochondrial metabolism may help correct post-COVID deficits in T cell memory responses.

Taken together, we have shown that even mild SARS-CoV-2 infection has lasting effects on T cell memory responses to other pathogens by limiting catabolic pathways such as glycolysis and fatty acid oxidation. This may explain why infection rates have increased in community and hospital settings post-pandemic and is of clear significance to the clinical management of infections in the post-pandemic era.

## Discussion

Post-pandemic, the US and the world at large have entered a “new normal” where morbidity and mortality rates across diseases have increased. Recent data from the UK revealed an excess death rate of 7.2% in 2022 and 8.6% in 2023 above expectations, despite the peak of the COVID-19 pandemic having passed^30^. Much of the increase in mortality can be attributed to higher rates of infections^31^, which raises the following question: *does COVID-19 affect host susceptibility to other infections?* This is the fundamental issue concerning this study.

Effective prevention and treatment of infectious diseases represent one of the seminal advances of medical science. From the discovery of germ theory to development of antibiotics and antivirals, inhabitants of the western world have become accustomed to mild infections with few long-term consequences over the past 80 years. The COVID-19 pandemic, however, reminded us that some infections remain deadly and have health consequences both during and beyond the acute infection stage. Here, we demonstrate that infection rates from additional pathogens across the US have increased in the years immediately following the introduction of COVID-19, which may be linked to immunometabolic dysfunction. We used matched PBMC samples from the same individuals before and after they contracted COVID-19 to determine how infection affected T cell memory responses to commonly encountered childhood and community-acquired pathogens. We found that post-COVID memory T cells have a unique transcriptional signature enriched in mitochondrial respiration pathways but deficient in T cell activation and MHC-II antigen presentation. Further, a majority of individuals exhibited post-COVID reductions of mitochondrial flux and ROS production in antigen-specific memory T cells, diminished connectivity of metabolic pathways (glycolysis, FAO, OXPHOS) in memory T cell populations, and an inability for antigen-specific T cells to mobilize catabolic pathways (glycolysis, FAO) after antigen stimulation (model in Fig. 6). These deficits were partially rescued by exposure to mitochondrial modulator drugs which may represent novel methods for immune reconstitution after COVID-19.

**Figure 6:**
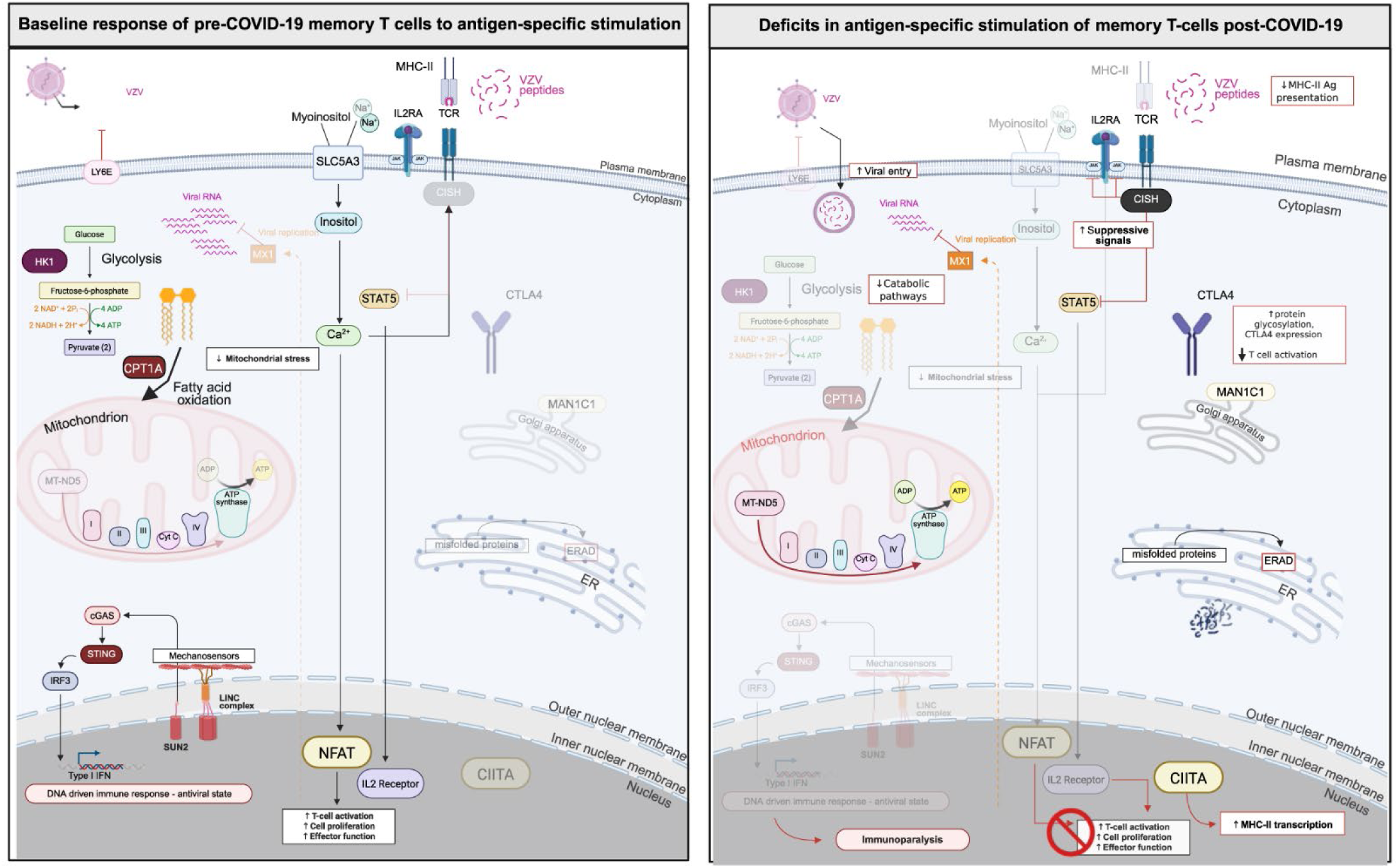
Post-COVID remodeling of antigen-specific T cell activation driven by metabolic reprogramming. Potential mechanisms of disrupted antigen-specific memory T cell activation post-COVID. Downregulated pathways are greyed out in each panel.

Emerging epidemiologic data underscore the broader clinical consequences of post-COVID immune remodeling. Multiple surveillance efforts have documented a sustained rise in infection rates across bacterial and viral pathogens in the post-pandemic period^32,33^. Importantly, this may be a unique consequence of COVID-19: patients with a positive COVID test between 2021 and 2023 had a significantly higher likelihood to contract bacterial bloodstream and urinary tract infections as well as herpes and respiratory viral infections compared with those seen for influenza A infection^32^. Mechanistically, T cell exhaustion and mitochondrial dysfunction have also been implicated in the pathogenesis of long COVID, where oxidative stress, dysregulated interferon signaling, and persistent immune activation converge to promote durable immunosuppression^13,14^. These converging lines of evidence suggest that SARS-CoV-2–induced dysregulation of adaptive immunity may not only compromise individual memory responses but also increase susceptibility to future outbreaks, including those of vaccine-preventable diseases.

The recall capacity of memory T cells is highly dependent on mitochondrial agility-i.e. the ability to switch between slow but efficient ATP generation with OXPHOS to quick but inefficient ATP generation with glycolysis and to an extent, fatty acid oxidation^34^. We found a unique post-COVID transcriptional signature in antigen-stimulated memory T cells with upregulation of immunosuppressive pathways and genes linked to mitochondrial respiration. The immunosuppressive genes *MAN1C1* which increases CTLA4 expression^35^ and *CISH*, an active suppressor of TCR signaling^36^, were also upregulated post-COVID while the myo-inositol transporter gene *SLC5A3* linked to NF-κB activation and T cell stimulatory gene *IL2Rb* were downregulated after infection. The *MX1* gene also remained upregulated in post-COVID samples despite being a marker of early and active SARS-CoV-2 infection which declined with viral clearance in patients enrolled in a human challenge study^37^. Given the considerable evidence that SARS-CoV-2 can persist in multiple tissues for months to years after infection^38^, it is possible that viral persistence plays a role in post-COVID T cell dysfunction, though we were unable to detect SARS-CoV-2 in PBMCs from our post-COVID samples (data not shown). Perhaps most importantly, mitochondrial oxidoreductase and respirasome genes were significantly upregulated after infection, suggesting that mitochondrial metabolism is disrupted in antigen-stimulated memory T cells after COVID-19 exposure which could be responsible for post-COVID immunosuppression.

While COVID infection did not alter the percentages of antigen-specific memory T cells, mitochondrial ROS production and electron flux across the mitochondrial membrane were significantly diminished. A closer examination of cell metabolism revealed a loss of connectivity between metabolic pathways in CD4 memory T cells after COVID-19. Mitochondrial dysfunction has routinely been identified within immune cells of long COVID patients^39^ and in COVID convalescents more broadly^40^, but we are the first to our knowledge to discover underlying mitochondrial dysfunction in antigen-specific memory T cells from patients *without* long COVID at 3-9 months after infection. It is possible that post-COVID immune suppression is a common feature of *any* COVID-19 infection, regardless of persistent symptoms. The extent to which COVID-induced immune suppression impacts susceptibility to other infections, particularly in vulnerable populations such as those with chronic disease or undergoing surgery, warrants further study.

Importantly, we found that glycolysis-related enzymes were upregulated in memory T cells at the population level but downregulated in antigen-specific memory T cells post-COVID. This validates our initial hypothesis that COVID-19 impairs the ability of resting memory T cells to switch from using OXPHOS to using glycolysis and FAO upon antigen re-encounter, which is critical to mount effective memory responses^25^. We think that the discordance between increased glycolysis usage in memory T cells at the population level but decreased glycolysis usage in antigen specific memory T cells represents a compensatory mechanism by which non-antigen specific T cells attempt to rectify the glycolytic deficit present in antigen specific cells. In a broader sense, this may mean that prior COVID-19 infection can prevent the antigen-specific memory cell from turning on catabolic pathways (glycolysis, FAO) to generate energy which may leave individuals more vulnerable to other infections in the medium to long-term. One intriguing possibility is that SARS-CoV-2 viral persistence may lead to chronic antigen stimulation and T cell exhaustion. Numerous studies have found SARS-CoV-2 viral persistence in tissues from the lung to the heart, lymph nodes, intestine, and even vasculature up to 2 years after COVID-19 infection^38,41^ and persistent antigen stimulation creates a state of T cell exhaustion in other chronic viral infections^42^. Though we did not detect SARS-CoV-2 viral RNA or protein in post-COVID PBMCs (Fig. S8), the impact of SARS-CoV-2 viral persistence on T cell memory warrants further study.

The study of post-COVID immune dysfunction has largely consisted of defining the problem observationally and mechanistically, and this includes studies from our group. Though elucidating the mechanism is critical to advancing and understanding of the problem, patients who currently suffer from immune suppression or other post-COVID sequelae often struggle to find effective treatment options. Here, we demonstrate that exposure to commonly available drugs that modulate mitochondrial function, metformin and ubiquinol, can partially correct post-COVID deficits in memory T cell metabolism. Metformin increases T cell function in numerous ways: it can decrease T cell senescence^43^, improve T cell fitness by reducing hypoxic damage^44^, and reprogram T cell metabolism to promote tumor clearance^45^. The mitochondrial complex III activator ubiquinol is a potent antioxidant which limits damage from ROS to promote T cell survival^46^. Perhaps most importantly, both are inexpensive commonly available drugs which may help to rectify post-COVID suppression of T cell memory responses.

Overall, this study showcases the durability of post-COVID immunometabolic remodeling in memory T cells in otherwise healthy convalescents. It will be critical to evaluate the extent to which SARS-CoV-2 viral persistence may contribute to memory T cell dysfunction in larger cohorts, particularly in vulnerable patient populations. Our data also suggest that clinical trials which repurpose commonly available and inexpensive drugs that promote mitochondrial function should be undertaken to combat post-COVID immune suppression by modulating immune metabolism. Finally, we think this study represents a step forward in understanding an emergent but underappreciated health crisis in post-COVID immune suppression.

### Limitations of study

The major limitation of this study is its sample size. Of the original 31 patients we enrolled, we were only able to collect data from subsets of the group for each figure due to sample availability constraints (ex. limited numbers of PBMCs isolated, study subjects contracted COVID-19 at different times). This made it difficult to include all samples in all experiments, though we strove to increase our sample size as much as possible. In particular, our sample size comes down to our decision to include matched samples only from individuals who were initially COVID-naïve, or unexposed. As the pandemic went on, identifying and enrolling unexposed individuals became more difficult and is almost impossible in the current day. We think future studies can build on our findings with larger sample sizes if they are not limited to COVID-naïve patients.

## Data availability

The data supporting the findings of this study are presented within the manuscript. RNA-Seq data will be deposited in the NCBI GEO database. All other data will be made available upon request.

## Code availability

All code and materials in the analysis of RNA Seq data are available via GitHub for the purpose of reproducing or extending the analysis. Original code for Boltzmann Brain analysis is proprietary and is unavailable due to imminent commercialization.

## Acknowledgments

We acknowledge the Northwestern University Genomics Core Facility and Illumina for providing free RNA sequencing (Illumina Pilot Grant to LV). We also acknowledge the Northwestern University and University of Chicago flow cytometry cores for their help in data gathering and analysis. K.C. acknowledges support from the German Research Foundation (DFG) through the Walter Benjamin Program [CI 469/2-1; 552519942].

## Author contributions

Conceptualization: L.V.; Investigation: D.C., J.A., J.S., L.L, N.R., S.Y., and L.V. Formal analysis: D.C., K.C., J.A., U.P., J.S, D.T., A.R., and L.V. Writing: K.C., M.W., J.C.A., and L.V., with input from all authors; Funding acquisition: L.V. Supervision: L.V.; Project administration: L.V.

## Methods

### Ethics Statement

Ethical approval for this study was obtained from the Northwestern University Institutional Review Board. All research on human subjects was conducted in accordance with the principles outlined in the Declaration of Helsinki and in accordance with institutional statutory requirements. All participants gave formal written consent to participate in the study and for their anonymized data to be published.

### Study Participant Information

We enrolled 31 healthy adult volunteers from Chicago, with an average age of 35. All participants provided informed consent and were recruited under IRB# STU00212583 from Northwestern University between 2021-2024. Subjects were enrolled after receiving COVID-19 mRNA vaccination and confirmed negative for SARS-CoV-2 by assessing antibody titers for SARS-CoV-2 nucleocapsid (figure 1C). Participants were vaccinated with the primary series of either the Pfizer BNT162B2 or Moderna mRNA-1273 mRNA vaccines during the course of the study. Blood samples were collected every two weeks during the first month, then monthly for 6 months. This sampling schedule was repeated following booster vaccination. Subjects were also instructed to report COVID-19 infection – following a positive PCR or antigen test – at any point within a year of vaccination. This approach enabled us to collect matched blood samples before and after infection, which were used for downstream assays following PBMC isolation. Additional demographic information can be found in Figure 1D. Comorbidities were self-reported and diagnosed prior to SARS-CoV-2 infection/vaccination.

### PBMC and Plasma Collection

Peripheral blood (30 mL) was collected from study participants both before and after SARS-CoV-2 infection. Samples were processed using a standard Ficoll density gradient protocol. Blood was diluted 1:1 with sterile 1x PBS and carefully layered (up to 30 mL) over 15 mL of Histopaque-1077 in 50 mL conical tubes. Tubes were centrifuged at 200 g for 30 minutes at room temperature. The upper plasma layer was aspirated and discarded, and the PBMC layer was carefully isolated without disturbing the RBC layer. Cells were cryopreserved in FBS + 10% DMSO at a concentration of 10–20 million cells/mL. Samples were frozen at –80°C and transferred to liquid nitrogen for long-term storage.

### Peptide Antigens used in cell stimulation

Peptide arrays for antigen stimulation were obtained from BEI Resources or from GenScript. Three arrays were used: Influenza Virus A (H1N1) Neuraminidase Protein, VZV Orf4 Protein, and *Staphylococcus aureus*-derived Hemolysin A. The Influenza virus peptide array consisted of 13-17mers with 11aa overlaps spanning the neuraminidase (NA) protein (BEI Resources). The VZV Orf4 peptide array consisted of 20mers with 10aa overlaps. Finally, the Hemolysin A peptide array consisted of peptides between 15 and 16mers with 10aa (both from GenScript). All three antigens were chosen for their ability to induce potent CD4 T-cell responses^47–49^.

### Antibody ELISA

Antigen-specific antibody titers were measured by ELISA as described previously^15^. In brief, 96-well flat-bottom MaxiSorp plates (Thermo Scientific) were coated with 1 µg/mL of one of the following recombinant proteins: SARS-CoV-2 Nucleocapsid, Influenza A Hemagglutinin, or hemolysin A from Staphylococcus aureus. Plates were then incubated at 4 °C for 48 hours and washed three times with wash buffer (PBS + 0.05% Tween 20). Blocking was performed with blocking solution (PBS + 0.05% Tween 20 + 2% bovine serum albumin), for 4 hr at room temperature. 6 µl of sera was added to 144 µl of blocking solution in the first column of the plate, 1:3 serial dilutions were performed until row 12 for each sample, and plates were incubated for 60 min at room temperature. Plates were washed three times with wash buffer followed by addition of secondary antibody conjugated to horseradish peroxidase, goat anti-human IgG (H + L) (Jackson ImmunoResearch) diluted in blocking solution (1:1000) and 100 µl/well was added and incubated for 60 min at room temperature. After washing plates three times with wash buffer, 100 µl/well of Sure Blue substrate (SeraCare) was added for 1 min. Reaction was stopped using 100 µl/well of KPL TMB Stop Solution (SeraCare). Absorbance was measured at 450 nm using a Spectramax Plus 384 (Molecular Devices). SARS-CoV-2 Nucleocapsid protein produced at the Northwestern Recombinant Protein Production Core by Dr. Sergii Pshenychnyi using plasmids that were produced under HHSN272201400008C and obtained from BEI Resources, NIAID, NIH: Vector pCAGGS containing the SARS-related coronavirus 2, NR-52309, nucleocapsid gene NR-53507. Purified H1N1 Hemagglutinin protein was obtained from R&D Biosystems. Recombinant *S. aureus* hemolysin A was purchased from LS Bio. VZV glyocoprotein E ELISA was purchased from ACRO Biosystems and the assay conducted according to manufacturer’s protocol.

### PBMC stimulation, T cell purification, and bulk RNA-Seq

Matched pre– and post-COVID PBMC samples from 16 individuals were randomly chosen for bulk RNA-Seq experiments. Cells were stimulated for 24h with 2µg/mL VZV-Orf4 peptides before cell isolation. CD4+CD45RO+ memory T cells were isolated by negative selection using the CD4 memory T cell isolation kit from Miltenyi Biotec. Cells were lysed and RNA extracted using the RNEasy Mini Kit (Qiagen). Library preparation was performed using the Illumina Stranded Total RNA-Seq Library Prep and sequenced using the Illumina NovaSEQ X Plus (Northwestern University Genomics Core Facility). Data were analyzed using the SCALES method from the Raman Lab^19^.

### Flow cytometry

For assessment of mitochondrial function, PBMCs were stimulated with IAV, SA, and VZV antigens for 16-18h. Cells were then stained with a surface antibody cocktail consisting of the following markers: CD3-BUV395, CCR7-BUV737, CD27-BUV805, CD8-V510, CD4-BV711, CD45RA-BV570, CD45RO-BV605, ICOS-AF700, CD69-BB700, PD1-BV421, CD137-BUV661, CD134-BV650 (all from BD Biosciences), CD14-APCFire 780, KLRG1-APCFire 810, Zombie UV live-dead, and CXCR5-PE-CF594 (all from Biolegend). For mitochondrial dye staining, MitoSOX Green, TMRM, and Mitotracker Red were purchased from Thermo Fisher. Cells were washed after surface staining and resuspended in warm PBS before the addition of mitochondrial dyes at the appropriate concentrations. Cells were incubated for 30min at 37°C, 5%CO_2_ in the dark before being washed in PBS and run through flow cytometry. Samples were acquired on a BD FACS Symphony 5 laser spectral flow cytometer at the Northwestern University Flow Cytometry Core Facility.

For Met-Flow, our staining procedure was adapted from Ahl et al^24^. After a 20-hour incubation with antigens +/− metformin 10µM^50^ or ubiquinol 5µM^51^, PBMCs were washed once with FACS wash buffer. Each well was stained with Live/Dead Blue dye before sequential surface and intracellular staining. Surface antibodies used were: CD3 (BUV395), CD4 (BV711), CD8 (BV510), CD69 (BB700), CD56 (BV650), PD1 (BV421), CD45RA (BV570), CD45RO (BV605), CD14 (APC-H7), CD16 (BV786), CD137 (BV750; all from Biolegend), CCR7 (BUV737), CD27 (BUV805), HLA-DR (BV480; all from BD Biosciences). For intracellular staining, the cells were incubated with an antibody cocktail containing the following antibodies: ASS1 (Coralite 594), ATP5a (PE-Alexa647), pMTOR (PE-Cy7), CPT1A (AF350; all ThermoFisher), ACAC (AF647), PRDX2 (BUV615), HK1 (BUV661), TKT (V450; all from BD Biosciences), G6PD (PE), GLUT1 (AF790; Santa Cruz Biotechnology). Samples were acquired on a Cytek Aurora 5 laser spectral flow cytometer at the University of Chicago Antibody and Cytometry Core Facility.

### Quantitative PCR to detect SARS-CoV-2 in PBMCs

RNA from PBMCs was isolated after antigen stimulation using the QIAamp DSP Viral RNA Mini Kit (Qiagen; Cat no. / ID. 61904) and RNA concentrations and quality were measured using the NanoDrop 2000 (ThermoFisher Scientific). Specific components from the NEB Luna Probe one-step RT-qPCR kit (New England Biolabs; cat. # M3029S, NEB Luna Probe One-Step RT-qPCR 4X Mix with UDG), SARS-CoV-2 Neo Assay Kit (Qiagen; Cat no. / ID. 222115, SARS-CoV-2 Neo Assay 20x primer/probe mix), and QIAprep&amp™ Viral RNA UM Kit (cat no. 221413, 221415, and 221417, human sampling IC assay; Mat. No. 1125243 and Lot No. 178039830) were utilized to prepare the assay mix of all components except for < 1 μg (total RNA) of the RNA template. To mix samples, gentle pipetting and inverting were completed. 18 µL of assay mix was pipetted first into each well. Following, samples had less than 1 µg of total RNA added to each well for up to 2 µL of volume, 2 µL total of positive controls (Orf1a/Orf1b – ROX and N1/N2 – FAM) or nuclease-free water as the negative control for a total of 20 µL volume reaction. Samples were run in technical duplicates using the QuantStudio 3 Flex Real-Time PCR Systems (Thermo Fisher Scientific) in a 96-well plate with a total reaction volume of 20 µL. Data analysis was conducted using QuantStudio Design & Analysis 2.x Software and visualized in a table using Ct/Cq values.

### Boltzmann Brain analysis

The matrix input into the Boltzmann Brain constituted X rows defining A and Y columns defining B, where each feature was either the percentage of cells in the population expressing a specified protein marker, or the geometric mean fluorescence intensity (gMFI) of protein expression on a per-cell basis derived from flow cytometry data analysis. The Boltzmann Brain program then discovered a statistical embedding that maximized the information between the statistical structure of variation defined by the matrix and the output variable of “Group”. This embedding comprised eigenmodes 4, 6, and 8 in Fig. 4J; features that were significantly distinct between “Group“ (i.e. pre-COVID vs. post-COVID) were extracted and are shown in the blue and red distributions in **Figs. 4** and **5**. The statistical metric used to determine features distinguishing pre– and post-COVID was the Jensen-Shannon distance (information based) in Fig. 4K and the Wasserstein earth mover’s distance (geometry based) in Fig. 5E.

### Statistical analysis

All data were analyzed in either GraphPad Prism 10 or using R/Python. All flow cytometry data were analyzed using FlowJo v10.

## Supplementary Figures

**Figure S1:**
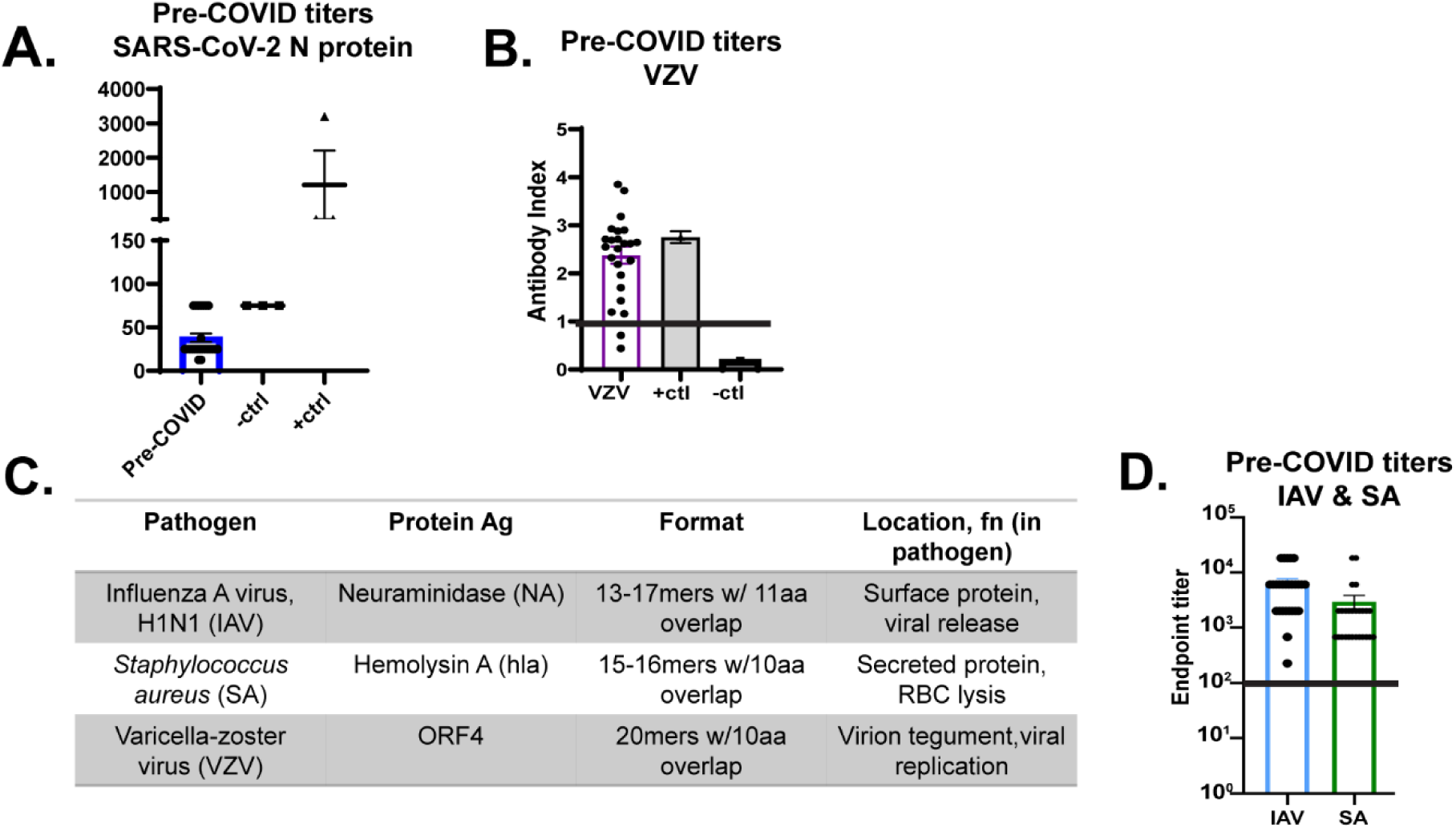
Pre-COVID antibody titers, VZV/SA/IAV antibody titers, antigen descriptions. A.) All subjects were negative for exposure to COVID-19 by anti-SARS-CoV-2 N protein (Nucleocapsid) serology. –ctrl: pre-2019 samples. +ctrl: patients diagnosed with long COVID after positive PCR test. B.) 93% of study subjects were positive for VZV antibody titers upon enrollment. All were confirmed to have been vaccinated or exposed to natural infection in childhood. –ctrl and +ctrl were provided with the manufacturer’s kit (ACRO Biosystems). C.) Description of antigens used in stimulation from VZV, IAV, and SA. D.) All subjects were positive for IAV and SA IgG by serology. Horizontal lines in B & D represent limit of detection.

**Figure S2:**
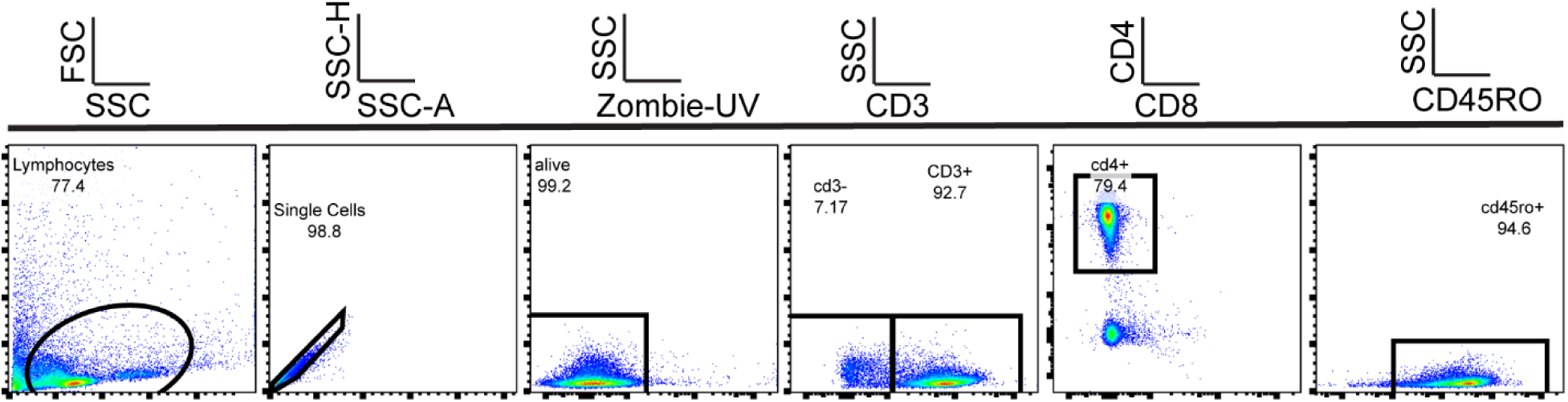
CD4 memory T cell purity after magnetic bead negative selection. PBMCs were stimulated with VZV-Orf4 peptides and CD4+CD45RO+ cells were purified by negative selection using a Miltenyi CD4 memory T cell purification kit. Samples were approximately 92% CD3+ T cells, of which 80% were CD4+.

**Figure S3:**
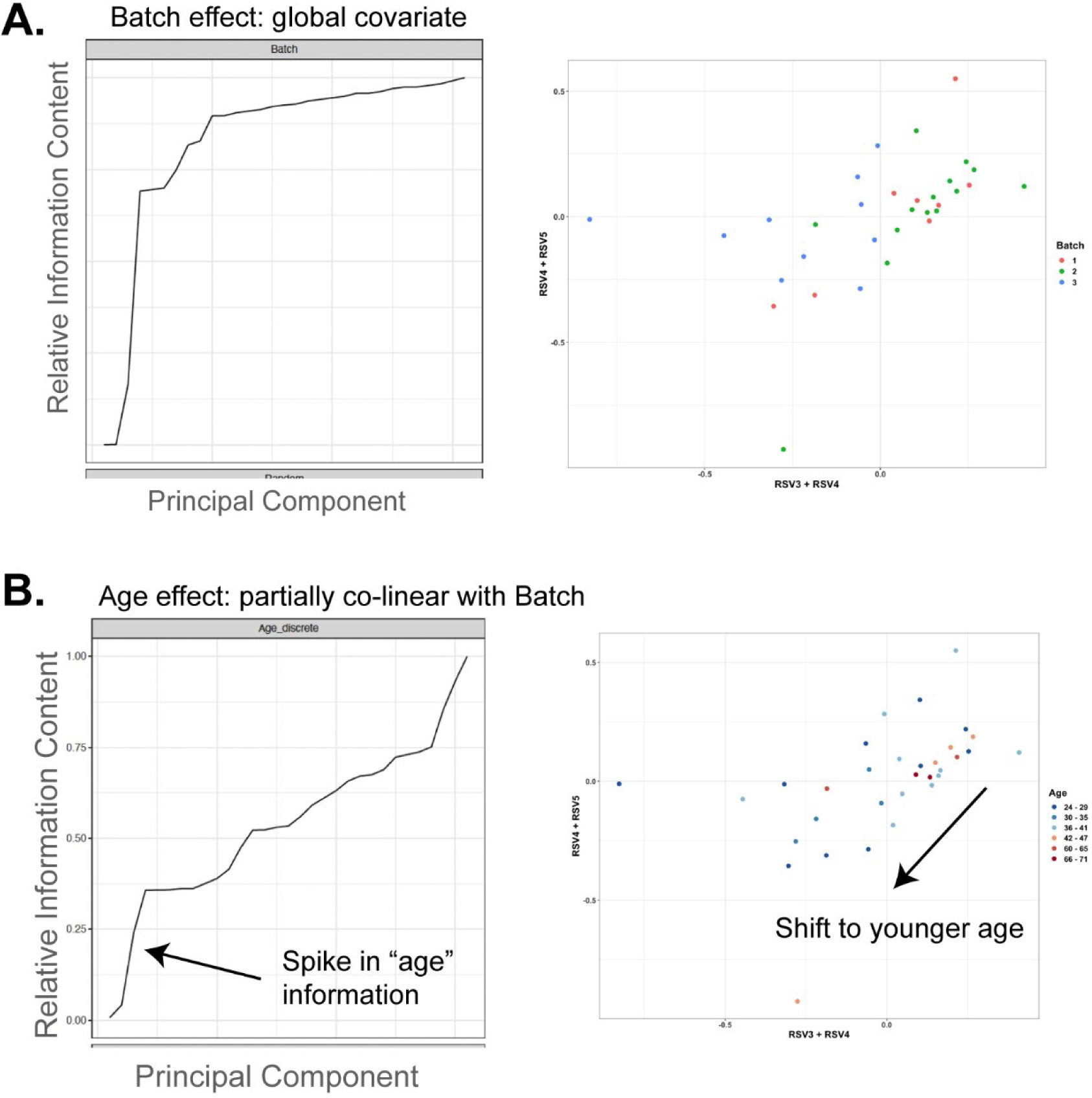
PCA analysis of pre– and post-COVID RNA-Seq data uncover batch and age effects as main global covariates. A.) Traditional PCA analysis identified batch effects as principal components describing statistical variation. B.) Age effects are similarly identified among the top PCs and are partially co-linear with batch effects.

**Figure S4:**
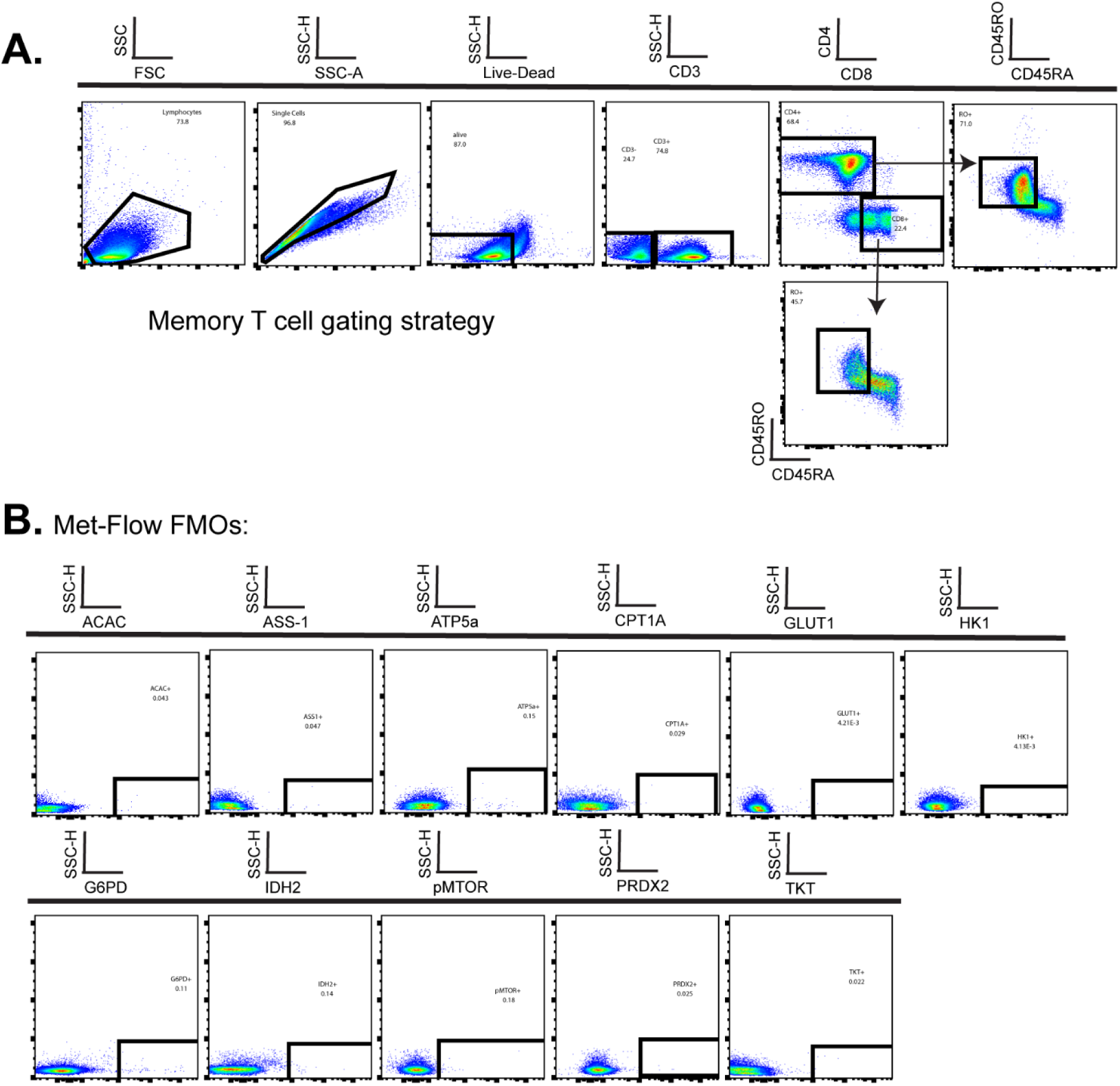
Flow cytometry gating strategies. A.) Strategy for gating CD4 and CD8 memory T cells. B.) Fluorescence minus one (FMO) gating for Met-Flow enzymes.

**Figure S5:**
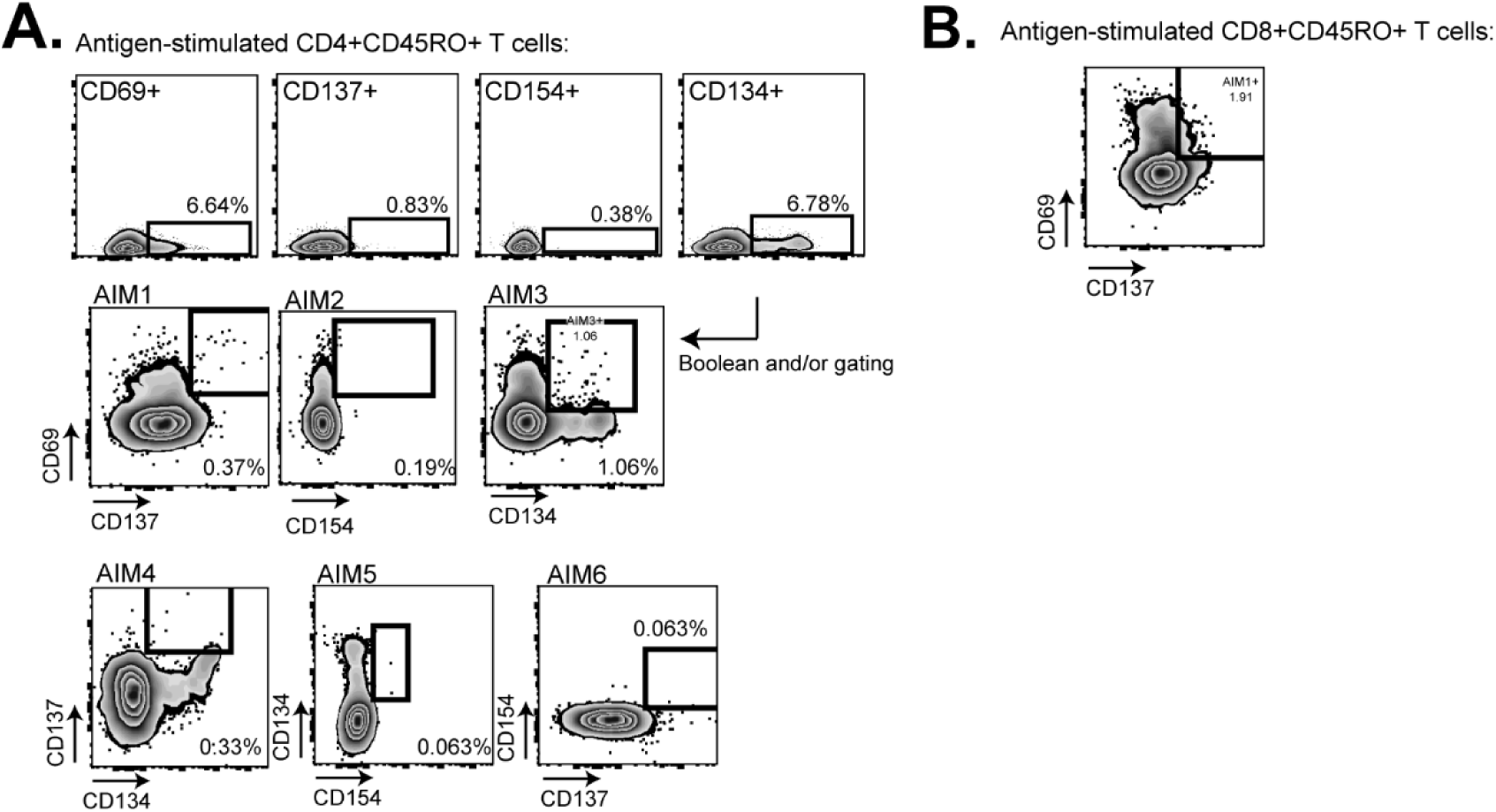
Gating strategy for AIM+ CD4 and CD8 memory T cells. A.) Boolean gating strategy was applied for identifying AIM+ CD4+CD45RO+ T cells. Cells were gated on CD69/CD137/CD154/CD134+ as in the top row before applying Boolean and/or gating (combination gates) in Flowjo. The 6 AIM combinations shown were concatenated together and metabolic enzyme expression analyzed within the concatenated population. B.) Gating strategy for AIM+CD8 memory T cells.

**Figure S6:**
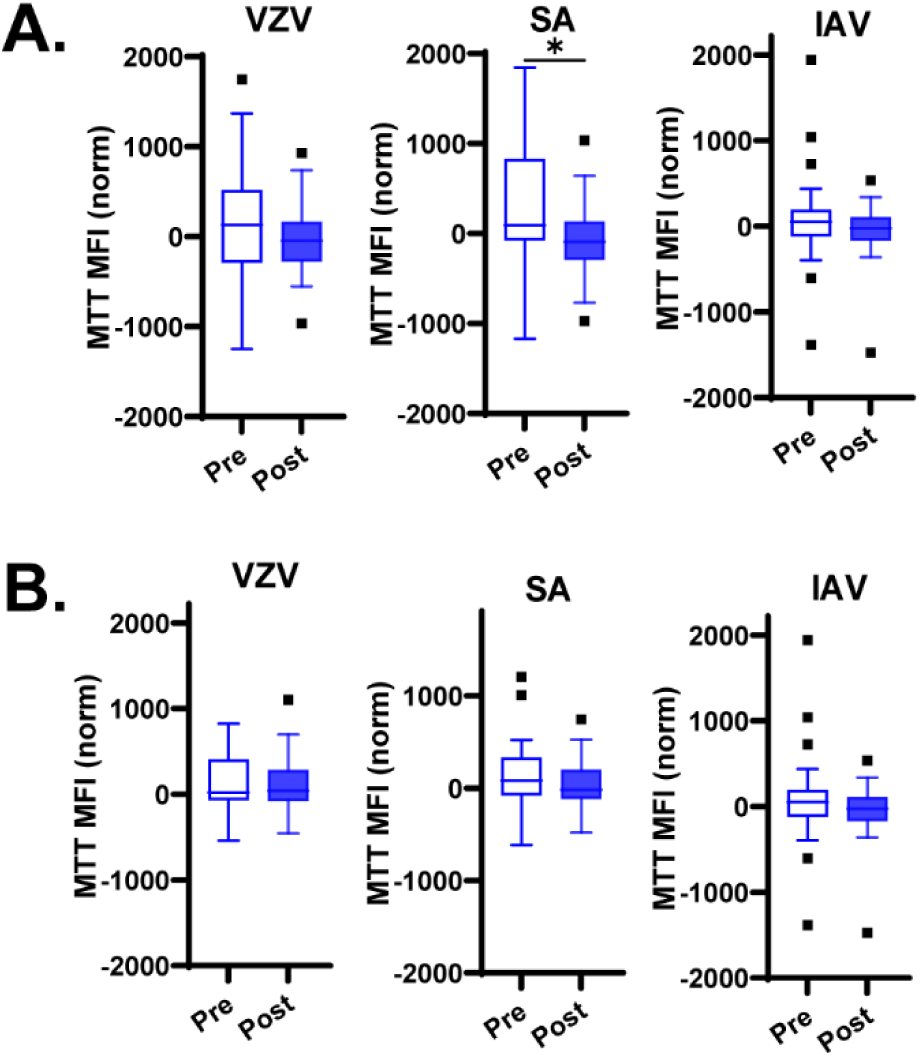
Active mitochondrial mass is similar in antigen-stimulated pre– and post-COVID AIM+ memory T cells. A.) Mitochondrial mass (MTT) is similar in pre– and post-COVID AIM+ CD4+CD45RO+ memory T cells, with the exception of SA stimulated T cells. B.) MTT is similar across all antigen stimulation conditions in AIM+CD8+CD45RO+ memory T cells.

**Figure S7:**
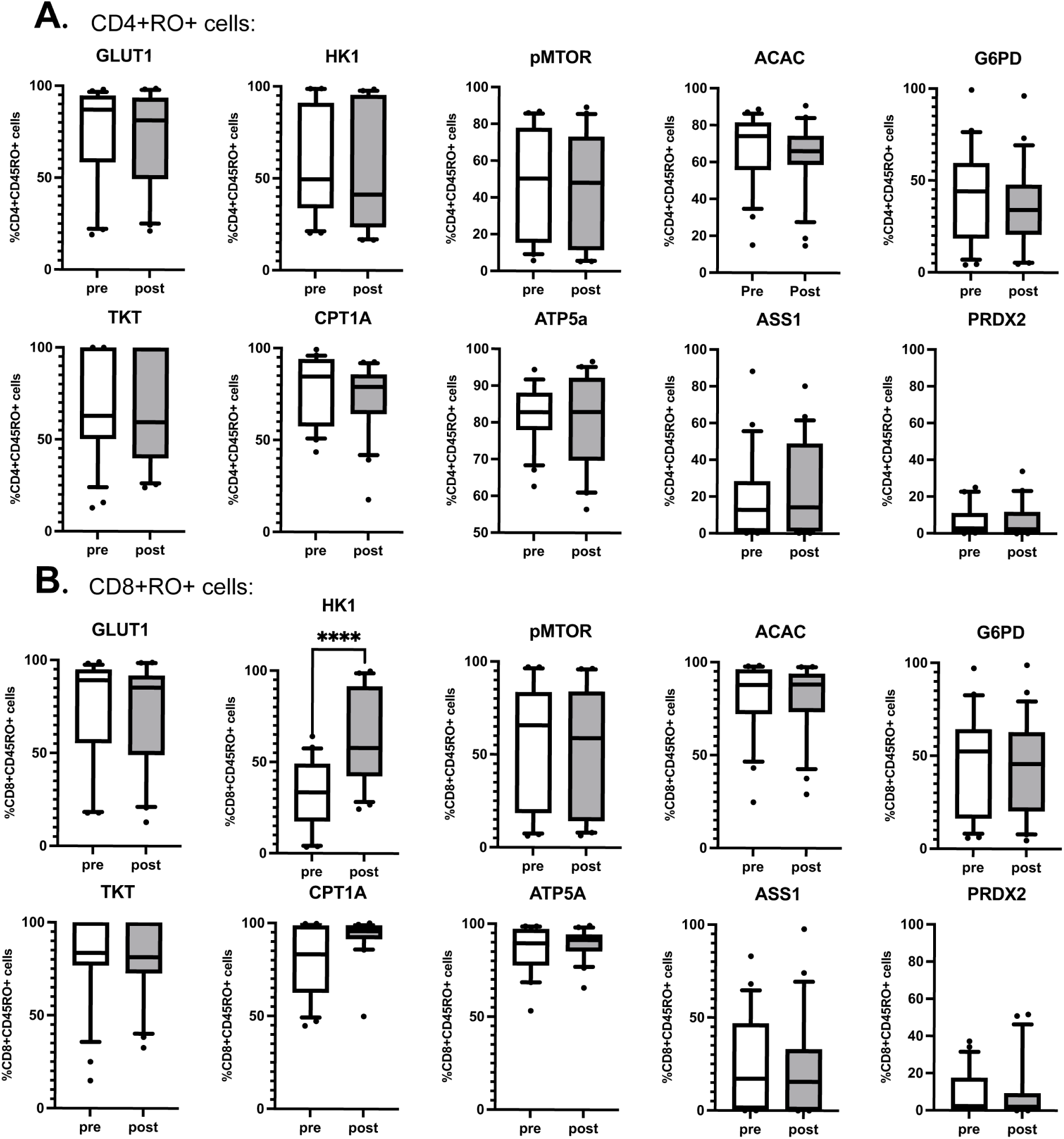
Met-Flow enzyme expression levels are similar in unstimulated pre– and post-COVID samples in memory T cells. A.) %CD4+CD45RO+ T cells positive for specified marker. B.) %CD8+CD45RO+ T cells positive for specified marker. ****p<0.001 by paired Student’s t test.

**Figure S8:**
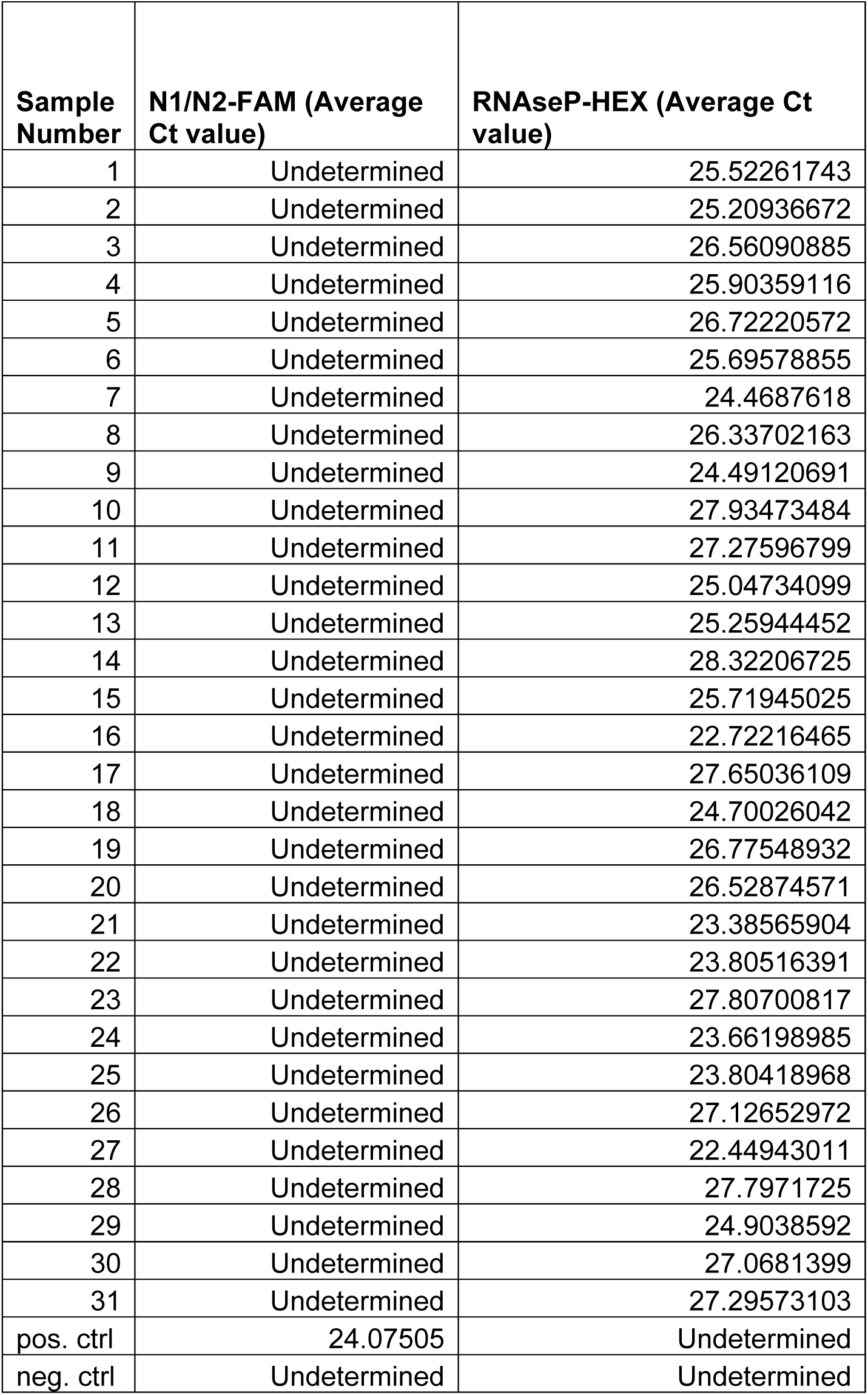
RT-qPCR cannot detect residual SARS-CoV-2 in post-COVID PBMCs. RT-qPCR detected RNA encoding SARS-CoV-2 Nucleocapsid protein within RNA isolated from post-COVID PBMCs.

## References

1. Lim, C. K. et al. Mpox diagnostics: Review of current and emerging technologies. J. Med. Virol. 95, e28429 (2023).

2. Luo, W. et al. Spatiotemporal variations of ‘triple-demic’ outbreaks of respiratory infections in the United States in the post-COVID-19 era. BMC Public Health 23, 2452 (2023).

3. Yek, C. et al. Impact of the COVID-19 Pandemic on Antibiotic Resistant Infection Burden in U.S. Hospitals: Retrospective Cohort Study of Trends and Risk Factors. Ann. Intern. Med. 178, 796–807 (2025).

4. Nickbakhsh, S. et al. Virus-virus interactions impact the population dynamics of influenza and the common cold. Proc. Natl. Acad. Sci. U. S. A. 116, 27142–27150 (2019).

5. Boehm, A. B. et al. More than a Tripledemic: Influenza A Virus, Respiratory Syncytial Virus, SARS-CoV-2, and Human Metapneumovirus in Wastewater during Winter 2022-2023. Environ. Sci. Technol. Lett. 10, 622–627 (2023).

6. Katz, J., Yue, S. & Xue, W. Herpes simplex and herpes zoster viruses in COVID-19 patients. Ir. J. Med. Sci. 191, 1093–1097 (2022).

7. O’Toole, R. F. The interface between COVID-19 and bacterial healthcare-associated infections. Clin. Microbiol. Infect. Off. Publ. Eur. Soc. Clin. Microbiol. Infect. Dis. 27, 1772–1776 (2021).

8. Clancy, C. J., Buehrle, D. J. & Nguyen, M. H. PRO: The COVID-19 pandemic will result in increased antimicrobial resistance rates. JAC-Antimicrob. Resist. 2, dlaa049 (2020).

9. Yek, C. et al. Impact of the COVID-19 Pandemic on Antibiotic Resistant Infection Burden in U.S. Hospitals: Retrospective Cohort Study of Trends and Risk Factors. Ann. Intern. Med. 178, 796–807 (2025).

10. Thompson, E. A. et al. Metabolic programs define dysfunctional immune responses in severe COVID-19 patients. Cell Rep. 34, 108863 (2021).

11. Siska, P. J. et al. Metabolic imbalance of T cells in COVID-19 is hallmarked by basigin and mitigated by dexamethasone. J. Clin. Invest. 131, e148225 (2021).

12. Geltink, R. I. K., Kyle, R. L. & Pearce, E. L. Unraveling the Complex Interplay Between T Cell Metabolism and Function. Annu. Rev. Immunol. 36, 461–488 (2018).

13. van der Windt, G. J. W. et al. CD8 memory T cells have a bioenergetic advantage that underlies their rapid recall ability. Proc. Natl. Acad. Sci. U. S. A. 110, 14336–14341 (2013).

14. Nankani, A. et al. Investigating demographic and geographic disparities in malnutrition and gastrointestinal cancer mortality among older adults in the United States: A comprehensive longitudinal Centers for Disease Control and Prevention WONDER analysis 1999-2020. Nutr. Clin. Pract. Off. Publ. Am. Soc. Parenter. Enter. Nutr. https://doi.org/10.1002/ncp.70056 (2025) doi:10.1002/ncp.70056.

15. Visvabharathy, L. et al. Neuro-PASC is characterized by enhanced CD4+ and diminished CD8+ T cell responses to SARS-CoV-2 Nucleocapsid protein. Front. Immunol. 14, 1155770 (2023).

16. Madsen, H. B. et al. Mitochondrial dysfunction in acute and post-acute phases of COVID-19 and risk of non-communicable diseases. Npj Metab. Health Dis. 2, 36 (2024).

17. Eberhardt, C. S. et al. Persistence of Varicella-Zoster Virus-Specific Plasma Cells in Adult Human Bone Marrow following Childhood Vaccination. J. Virol. 94, e02127–19 (2020).

18. Lu, C.-L., Wang, J., Chang, Y.-C. & Lu, K.-C. Cardiorenal outcomes after herpes zoster reactivation in COVID-19 survivors from a global TriNetX study. Sci. Rep. 15, 30036 (2025).

19. Zaydman, M. A. et al. Defining hierarchical protein interaction networks from spectral analysis of bacterial proteomes. eLife 11, e74104 (2022).

20. Zhang, J., Lyu, T., Cao, Y. & Feng, H. Role of TCF-1 in differentiation, exhaustion, and memory of CD8+ T cells: A review. FASEB J. Off. Publ. Fed. Am. Soc. Exp. Biol. 35, e21549 (2021).

21. Ma, Y. et al. Glycosylphosphatidylinositol biosynthesis functions as a conserved host defense pathway against coronaviruses via regulation of LY6E. PLOS Pathog. 21, e1013441 (2025).

22. Torisu, H. et al. Functional MxA promoter polymorphism associated with subacute sclerosing panencephalitis. Neurology 62, 457–460 (2004).

23. Palmer, D. C. et al. Cish actively silences TCR signaling in CD8+ T cells to maintain tumor tolerance. J. Exp. Med. 212, 2095–2113 (2015).

24. Ahl, P. J. et al. Met-Flow, a strategy for single-cell metabolic analysis highlights dynamic changes in immune subpopulations. *Commun*. Biol. 3, 305 (2020).

25. Pearce, E. L. et al. Enhancing CD8 T-cell memory by modulating fatty acid metabolism. Nature 460, 103–107 (2009).

26. Yang, M. et al. Inhibition of mitochondrial function by metformin increases glucose uptake, glycolysis and GDF-15 release from intestinal cells. Sci. Rep. 11, 2529 (2021).

27. Baigorri, R. E. et al. Metformin improves CD8+ T cell responses and parasitemia control via macrophage modulation during Trypanosoma cruzi infection. J. Leukoc. Biol. qiaf134 (2025) doi:10.1093/jleuko/qiaf134.

28. Zhang, Z. et al. Metformin Enhances the Antitumor Activity of CD8+ T Lymphocytes via the AMPK-miR-107-Eomes-PD-1 Pathway. J. Immunol. Baltim. Md 1950 204, 2575–2588 (2020).

29. Reina-Campos, M. et al. Metabolic programs of T cell tissue residency empower tumour immunity. Nature 621, 179–187 (2023).

30. Pearson-Stuttard, J., Caul, S., McDonald, S., Whamond, E. & Newton, J. N. Excess mortality in England post COVID-19 pandemic: implications for secondary prevention. Lancet Reg. Health Eur. 36, 100802 (2024).

31. Feinmann, J. Analysis reveals global post-covid surge in infectious diseases. BMJ 385, q1348 (2024).

32. Cai, M., Xu, E., Xie, Y. & Al-Aly, Z. Rates of infection with other pathogens after a positive COVID-19 test versus a negative test in US veterans (November, 2021, to December, 2023): a retrospective cohort study. Lancet Infect. Dis. 25, 847–860 (2025).

33. Biggs, H. M. et al. Trends in Incidence and Epidemiology of Methicillin-Resistant Staphylococcus aureus Bacteremia, Six Emerging Infections Program Surveillance Sites, 2005-2022. Open Forum Infect. Dis. 12, ofaf282 (2025).

34. Corrado, M. & Pearce, E. L. Targeting memory T cell metabolism to improve immunity. J. Clin. Invest. 132, (2022).

35. Chen, H.-L., Li, C. F., Grigorian, A., Tian, W. & Demetriou, M. T Cell Receptor Signaling Co-regulates Multiple Golgi Genes to Enhance N-Glycan Branching♦. J. Biol. Chem. 284, 32454–32461 (2009).

36. Palmer, D. C. et al. Cish actively silences TCR signaling in CD8+ T cells to maintain tumor tolerance. J. Exp. Med. 212, 2095–2113 (2015).

37. Rosenheim, J. et al. SARS-CoV-2 human challenge reveals biomarkers that discriminate early and late phases of respiratory viral infections. Nat. Commun. 15, 10434 (2024).

38. Stein, S. R. et al. SARS-CoV-2 infection and persistence in the human body and brain at autopsy. Nature 612, 758–763 (2022).

39. Maison, D. P. et al. Peripheral immune progression to long COVID is associated with mitochondrial gene transcription: A meta-analysis. Mitochondrion 85, 102072 (2025).

40. Lage, S. L. et al. Persistent immune dysregulation and metabolic alterations following SARS-CoV-2 infection. MedRxiv Prepr. Serv. Health Sci. 2025.04.16.25325949 (2025) doi:10.1101/2025.04.16.25325949.

41. Peluso, M. J. et al. Tissue-based T cell activation and viral RNA persist for up to 2 years after SARS-CoV-2 infection. Sci. Transl. Med. 16, eadk3295 (2024).

42. Kahan, S. M., Wherry, E. J. & Zajac, A. J. T cell exhaustion during persistent viral infections. Virology 479-480, 180–193 (2015).

43. Yang, J. et al. The effect of metformin on senescence of T lymphocytes. Immun. Ageing A 20, 73 (2023).

44. Finisguerra, V. et al. Metformin improves cancer immunotherapy by directly rescuing tumor-infiltrating CD8 T lymphocytes from hypoxia-induced immunosuppression. J. Immunother. Cancer 11, e005719 (2023).

45. Huang, X. et al. Metformin Reprograms Tryptophan Metabolism to Stimulate CD8+ T-cell Function in Colorectal Cancer. Cancer Res. 83, 2358–2371 (2023).

46. Gollapudi, S. & Gupta, S. Reversal of oxidative stress-induced apoptosis in T and B lymphocytes by Coenzyme Q10 (CoQ10). Am. J. Clin. Exp. Immunol. 5, 41–47 (2016).

47. Milikan, J. C. M. et al. Identification of viral antigens recognized by ocular infiltrating T cells from patients with varicella zoster virus-induced uveitis. Invest. Ophthalmol. Vis. Sci. 48, 3689–3697 (2007).

48. Babon, J. A. B. et al. Genome-wide screening of human T-cell epitopes in influenza A virus reveals a broad spectrum of CD4+ T-cell responses to internal proteins, hemagglutinins, and neuraminidases. Hum. Immunol. 70, 711–721 (2009).

49. Wang, K. et al. Alpha hemolysin enhances the immune response by modulating dendritic cell differentiation via ADAM10-Notch signaling. Signal Transduct. Target. Ther. 10, 334 (2025).

50. Chao, R. et al. Nutrient Condition in the Microenvironment Determines Essential Metabolisms of CD8+ T Cells for Enhanced IFNγ Production by Metformin. Front. Immunol. 13, 864225 (2022).

51. Okamoto, T. et al. Protective effect of coenzyme Q10 on cultured skeletal muscle cell injury induced by continuous electric field stimulation. Biochem. Biophys. Res. Commun. 216, 1006–1012 (1995).

